# BCL6-dependent TCF-1^+^ progenitor cells maintain effector and helper CD4 T cell responses to persistent antigen

**DOI:** 10.1101/2021.08.06.455141

**Authors:** Yu Xia, Katalin Sandor, Joy A. Pai, Bence Daniel, Saravanan Raju, Renee Wu, Sunnie Hsiung, Yanyan Qi, Tenzin Yangdon, Mariko Okamoto, Robert D. Schreiber, Kenneth M. Murphy, Ansuman T. Satpathy, Takeshi Egawa

## Abstract

Shortly after priming, the fate of activated CD4 T cells is segregated into BCL6^+^ follicular helper T (Tfh) and BCL6^−^ effector (Teff) cells. However, it remains unknown how these subsets are sustained in the presence of chronic antigen stimulation. Using a combination of single cell- and population-based approaches, we show that in chronic viral infection, activated CD4 T cells differentiate into BCL6-dependent TCF-1^+^ progenitor cells with superior capacity to expand and give rise to both Teff and Tfh. They share properties with progenitor-exhausted CD8 T cells and are required for the continued generation of Teff cells as antigen persists. In response to tumors, an analogous CD4 T cell population develops in draining lymph nodes. Our study reveals the heterogeneity and plasticity of CD4 T cells upon encountering persistent antigen and highlights their population dynamics through a stable bipotent intermediate state.

## Introduction

CD4 T cells play a central role in coordinating innate and adaptive immunity. In response to signals through T cell receptors (TCR) and cytokines provided by dendritic cells, they functionally differentiate into T effector (Teff) cells and produce context-specific cytokines to coordinate activation of innate immune cells (Crotty et al., 2010; Marshall et al., 2011; Pepper and Jenkins, 2011). Activated CD4 T cells also differentiate into follicular helper T (Tfh) cells that provide contact-dependent help to B cells for their expansion and affinity maturation. Shortly after priming, CD4 T cells diverge into BCL6^−^ and BCL6^+^ subsets depending on the strength of TCR and IL-2R signaling in a B cell-independent manner (Baumjohann et al., 2011; Choi et al., 2011; DiToro et al., 2018; Hondowicz et al., 2018). Stronger TCR signaling and subsequent upregulation of IL-2 is associated with the expression of BCL6 and differentiation into Tfh cells whereas activation of IL-2R signaling and IL-12R signaling induces upregulation of the transcriptionation factor Blimp1, which promotes Teff differentiation and suppresses Tfh fate via downregulation of BCL6 and TCF-1 (DiToro et al., 2018; Johnston et al., 2009; Sheikh and Groom, 2021; Wu et al., 2015). In studies using acute pathogen infection or non-replicating antigen, the early binary fate decision of CD4 T cells is stable as early BCL6^+^ cells exhibit skewed developmental potential towards Tfh at the expense of Teff cells (Choi et al., 2013; DiToro et al., 2018).

Following antigen clearance, the antigen-specific CD4 T cell population contracts, giving rise to memory CD4 T cells that confer long-lasting protection against re-infection. These cells are classified into central (Tcm) and effector (Tem) memory subpopulations (Ruterbusch et al., 2020). Tem cells, similar to Teff cells, express tissue homing receptors, promptly produce effector cytokines such as interferon-gamma (IFN-g) upon rechallenge, and decay faster than Tcm. In contrast, Tcm cells express homing receptors to lymphoid organs, such as CCR7 and CD62L, and are capable of robust IL-2 production immediately after stimulation, resembling BCL6^+^ helper cells that emerge early after priming. In acute viral or intracellular bacterial infections, Tcm cells are derived from central memory precursor CD4 T cells (Tcmp) that are formed at the peak of the T cell response (Ciucci et al., 2019; Marshall et al., 2011; Pepper et al., 2011). Tcmp cells already express Tcm-related genes such as *Tcf7, Ccr7, Id3* and *Bcl2*, but phenotypically overlap with pre-Tfh cells with an intermediate level of BCL6 and CXCR5 expression (Crotty, 2011, 2019). Thus, the relationship between Tfh and memory populations and the nature of memory precursor cells has remained incompletely understood.

CD4 T cells also play essential roles in controlling chronic viral infection and tumors, in which their responses must be sustained for substantially longer periods, compared to acute infection or vaccine-elicited immune responses. Similar to CD8 T cells, activated CD4 T cells acquire distinct phenotypes in the presence of chronic antigen, compared to those induced by acute viral infection (Crawford et al., 2014; Hashimoto et al., 2018; Wherry et al., 2007). Whereas increased expression of inhibitory receptors is the hallmark feature of exhausted CD8 T cells, CD4 T cells exposed to persistent antigen exhibit a unique pattern of gene expression changes, including elevated expression of *Bcl6* and additional similarities to Tfh cells (Fahey et al., 2011), while expression of multiple inhibitory receptors is less pronounced compared to CD8 T cells (Crawford et al., 2014). In CD8 T cells, a series of recent studies identified a subset of stem-like or progenitor-like exhausted CD8 T cells, referred to as T progenitor exhausted (TPEX) cells, that express the transcription factors (TF) TCF-1 and BCL6, and the canonical Tfh marker CXCR5 (He et al., 2016; Im et al., 2016; Leong et al., 2016; Utzschneider et al., 2016; Wu et al., 2016). TPEX cells are necessary to sustain CD8 T cell effector responses and critical for enhanced antiviral and anti-tumor immunity in response to immune checkpoint blockade. Given the terminally differentiated nature of CD4 Teff and Tfh cells, the continued CD4 T cell response by Teff and Tfh cells may similarly be supported by their continued differentiation from less differentiated stem-like, or progenitor, cells. However, it remains unknown whether sustained CD4 T cell response depends on such a progenitor population, or how they are distinct, if any, from pre-Tfh or memory CD4 T cells.

To address this question, we performed an unbiased characterization of the heterogeneity of antigen-specific CD4 T cells in response to infection with a chronic strain of *Lymphocytic choriomeningitis virus* (LCMV, clone 13 (c13)), which achieves prolonged antigen persistence, infection and identified a population of PD-1^+^ progenitor CD4 T (Tprog) cells that were TCF-1^+^ BCL6^lo/−^ and distinct from TCF-1^−^ BCL6^−^ Teff and TCF-1^+^ BCL6^hi^ Tfh cells. This cell population is detectable at the peak of T cell response and persists into the chronic phase of infection in a B cell-independent manner. Trajectory analysis, TCR clonal tracing, and adoptive transfer experiments, demonstrate that they serve as common progenitors for both Teff and Tfh cells. Epigenomic analysis with single-cell ATAC-seq (scATAC-seq) suggests that TCF-1^+^ BCL6^lo/−^ progenitor cells require NFAT and AP-1 TF activity for early differentiation, followed by bifurcation to the two distinct terminal fates. Their development also requires BCL6 and CD4 T cell-specific conditional deletion of *Bcl6* not only results in the expected loss of BCL6^hi^ TCF-1^+^ Tfh but also the total ablation of TCF-1^−^ BCL6^−^ Teff cells two weeks after infection, despite an intact initial expansion. Finally, in tumor-bearing mice, phenotypes of tumor-reactive CD4 T cells were distinct between the tumor microenvironment and tumor-draining lymph nodes (tdLNs). An analogous heterogeneity of CD4 T cells was observed in tdLNs, while antigen-specific CD4 T cells in the tumor were exclusively TCF-1^−^ Teff cells, suggesting tumor-specific CD4 T cells are maintained in secondary lymphoid organs rather than in the tumor microenvironment. These results collectively indicate that following an initial wave of naive T cell-derived differentiation of Teff and Tfh cells, CD4 Teff and Tfh cell responses to persistent antigen are maintained by a pool of BCL6-dependent TCF-1^+^ common Tprog cells, akin to how the CD8 response is maintained by CD8 TPEX.

## Results

### scRNA-seq and flow cytometry reveals heterogeneity within antigen-specific CD4 T cells responding to chronic viral infection

To obtain an unbiased overview of the CD4 T cell response to persisting antigen, we performed paired single-cell RNA-seq (scRNA-seq) and TCR sequencing of I-A^b^-LCMV-gp66-specific (Tet^+^) and the remaining total CD4 T cells (Tet^−^) in C57BL/6 (B6) mice infected either with LCMV-c13 (collected 8 or 21 days post infection - dpi) or with an acute strain of LCMV (Armstrong strain (Arm)) (collected 8 dpi). In total, we obtained 20,988 cells after quality control filtering based on the number of genes captured and percent mitochondrial reads per cell (Methods), and we obtained paired *TCRαβ* sequences in 95.6% of these cells (**Fig. S1A**). Dimensionality reduction and clustering of the cells based on their gene expression profiles identified 15 cell type clusters, most of which were grouped into 5 large clusters: *Sell*^+^ *Ccr7*^+^ *Tcf7*^+^ *Slamf6*^−^ naive (Cluster 1, 2 from RNA-seq as abbreviated as C1r, 2r), *Tcf7*^+^ *Slamf6*^+^ *Bcl6*^−^ *Cxcr5^lo/−^* resting memory-like (C3r, 4r), *Tcf7*^+^ *Slamf6*^+^ activated memory-like (C5r and 6r), *Tcf7*^+^ *Slamf6*^+^ *Bcl6*^+^ *Cxcr5*^+^ Tfh (C7r), and *Prdm1^+^ Slamf1^+^ Cxcr6*^+^ Teff cells (C8r-10r) (**Fig. 1A, B**). C5r and C6r were characterized by the expression of *Tcf7, Slamf6, Pdcd1*, and *Bcl2* at similar levels to C3r and C4r, and additionally expressed *Cd69* and *Mif*, which are upregulated by activated T cells (Choi et al., 2012; Ziegler et al., 1994). They also expressed additional genes induced by TCR signaling, including *Egr1, Egr2, Nr4a1, Tnfrsf4, Nfkbia* and *Batf* (**Fig. 1A-C; Fig. S1B, C**), suggesting that they represent activated memory-like cells. We also identified additional small clusters, including thymic *Il2ra*^hi^ and peripheral *Il2ra^lo^* regulatory T cells (Tregs; C11r and 12r, respectively), and an undefined population of non-Treg CD4 T cells with elevated IFN-stimulated genes and MYB expression (C13r and 14r, respectively) and the chemokine receptors, *Ccr4* and *Ccr6* (C15r).

**Figure 1.**
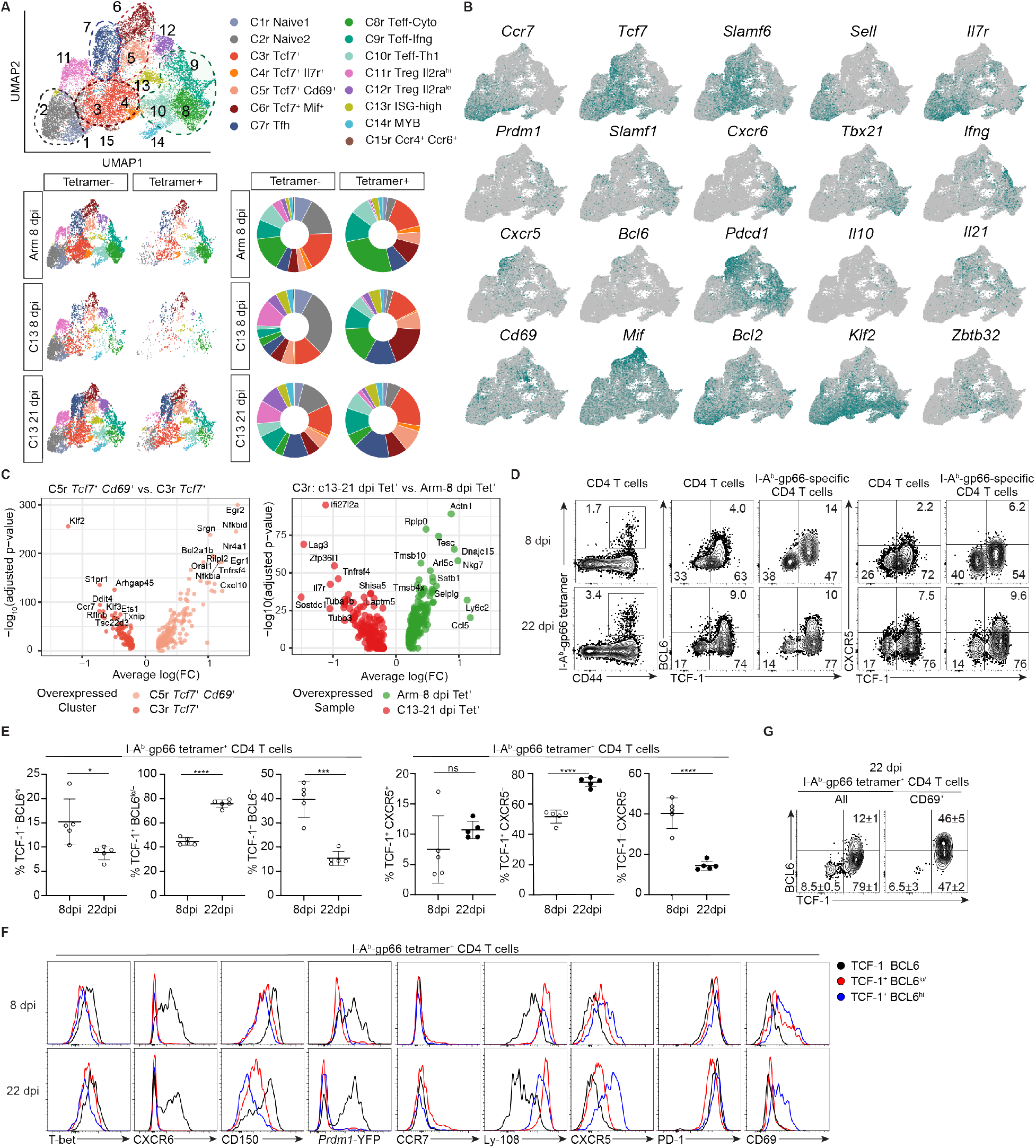
scRNA-seq and flow analysis reveals heterogeneity within antigen-specific CD4 T cells in chronic LCMV infections. **(A)** UMAP of LCMV-Arm d8, LCMV-c13 d8 and d21 CD4 T splenocytes sorted for I-Ab-gp66 Tet^+^ and Tet^−^ populations. Cells are clustered by scRNA-seq gene expression profiles and colored by phenotype (top). UMAP of CD4 T cells from each condition and tetramer-sorted population (bottom left). Proportion of cells from each condition in each phenotypic cluster (bottom right). **(B)** Expression of cluster-defining markers. **(C)** Differential gene expression of cells in the activated memory-like cluster C5r compared to the progenitor/memory cluster C3r (left). Differential gene expression of progenitor C3r cells from Arm d8 compared to cl13 d21 (right). **(D-E)** Expression of TCF-1, BCL6, and CXCR5 by I-Ab-gp66-specific splenic CD4 T cells in B6 mice infected with LCMV-c13 8 or 22 days before the analysis. Representative flow cytometry plots are shown with gating and frequencies of cells and pooled data from xx experiments (n = xx / experiment) are shown with mean±SD in (E) with unpaired *t*-test. **(F-G)** Expression of indicated genes in I-Ab-gp66 specific splenic CD4 T cell subpopulations harvested from LCMV-c13 infected B6 mice. Data are representative of xx experiments.

Arm-8 dpi Tet^−^ cells, which contained a mixture of bystander naive cells, resting memory CD4 T cells, and LCMV-reactive CD4 T cells recognizing non-dominant epitopes, were present in all 5 major clusters (**Fig. 1A**). In contrast, Tet^+^ cells predominantly exhibited Teff and Tfh phenotypes, constituting approximately 60% of the whole population. A large proportion of the remaining Tet^+^ cells displayed the resting memory phenotype (**Fig. 1A**), likely corresponding to Tcmp (Ciucci et al., 2019; Pepper et al., 2011). In c13 infection, the relative frequency of Teff cells in Tet^−^ CD4 T cells was significantly reduced compared to CD4 T cells responding to Arm infection, while that of Tfh cells was similar, indicating Tfh-skewing of expanding CD4 T cells (**Fig. 1A**). As in Arm infection, Teff cells were enriched in Tet^+^ cells compared to Tet^−^ cells. However, compared to Tet^+^ cells in Arm infection, the percentage of Teff cells within Tet^+^ cells in c13 infection was notably reduced on both 8 dpi and 21 dpi (**Fig. 1A**). This reduction of Teff cells was associated with an increased proportion of Tfh cells in c13 infection, especially at 21 dpi. Frequencies of Tet^+^ cells with the resting memory phenotype (C3r and 4r) were comparable between Arm and c13 infection on 8 dpi, and 8 dpi and 21 dpi in c13 infection. However, in c13-Tet^+^ samples, these clusters differed qualitatively from the Arm counterparts with a relative upregulation of activation-induced genes (e.g., *Lag3, Tnfrsf4*), ribosomal protein subunits, and IFN-I-induced genes (e.g., *Ifi27l2a*) which are associated with Tfh cells (Ciucci et al., 2019), and downregulation of the Th1/Teff-associated genes, *Ccl5, Nkg7* and *Ly6c2* (Jenner et al., 2009; Lund et al., 2005; Marshall et al., 2011)(**Fig. 1C, right**). These changes in gene expression suggest that cells in the memory-like cell populations are biased towards Tfh over Teff. In addition, activated memory-like cells in C5r and 6r were increased in c13-Tet^+^ samples compared to Arm infection samples, suggesting increased mobilization from the memory-like cell pool into more differentiated cells (**Fig. 1A**).

To validate the scRNA-seq dataset with analysis of the subset-specific proteins, we used flow cytometry to analyze splenocytes of c13-infected B6 mice on 8 and 22 dpi for BCL6 and TCF-1 expression (**Fig. 1D-F**). On 8 dpi, Tet^+^ CD4 T cells were largely separated into TCF-1^+^ and TCF-1^−^ cells (**Fig. 1D**). TCF-1^+^ cells uniformly expressed intermediate levels of BCL6 and CXCR5 proteins. On 22 dpi, TCF-1^+^ cells constituted a large proportion of Tet^+^ CD4 T cells, and a subfraction of these cells upregulated BCL6 and CXCR5 as they further underwent differentiation into Tfh cells, whereas the rest of TCF-1^+^ cells downregulated BCL6 to a level similar to that of TCF-1^−^ cells (**Fig. 1D, E**). Consistent with the scRNA-seq data, approximately 15% of Tet^+^ CD4 T cells expressed CD69, and about half of them were TCF1^+^ BCL6^lo/−^. TCF-1^−^ cells barely expressed CD69, consistent with the scRNA-seq data. The remaining half of CD69^+^ cells were TCF1^+^ BCL6^+^ Tfh cells, possibly reflecting preferential differentiation of cells that had received stronger TCR signals and expressed higher Cd69 to Tfh (**Fig. 1G**). Both TCF-1^+^ populations expressed Ly-108 (encoded by *Slamf6*) but lacked expression of the Teff marker *Prdm1* based on a *Prdm1-YFP* reporter (**Fig. 1F**). TCF-1^+^ BCL6^lo/−^ cells expressed lower PD-1 and higher Ly-108 compared to TCF-1^+^ BCL6^hi^ cells, and a minority (~5%) of them expressed CCR7, in contrast to resting Tcm. TCF-1^−^ cells upregulated T-bet, CXCR6 and CD150 (encoded by *Slamf1*) as well as the *Prdm1* reporter, indicating that they were highly enriched for differentiated Teff cells. TCF-1^−^ cells also contained a small percentage (~5%) of Foxp3^+^ Treg cells (**Fig. S1D, E**). Consistent with previous reports, CXCR5 was a reliable surrogate marker of BCL6, while Ly-108 was a reliable surrogate for TCF-1 expression (**Fig. S1F)**. In terms of tissue distribution, Tfh cells were barely detected in the bone marrow or liver, whereas Teff cells were detected at higher percentages (40-50%) in those organs compared to the spleen, and TCF-1^+^ BCL6^lo/−^ cells were also present in those organs (**Fig. S1G, H)**.

Collectively, our results indicate that with chronic persistence of antigen, a substantial fraction of antigen-specific CD4 T cells acquire a TCF-1^+^ *Slamf6^+^ Bcl2*^+^ PD-1^+^ memory-like phenotype. These memory-like CD4 T cells resemble CD8 TPEX and contain a subpopulation of cells with a gene expression signature associated with TCR-mediated activation. Additionally, their frequencies were increased in LCMV-c13 infection, indicative of progenitor potential, actively supporting the maintenance of Teff and Tfh cells in response to persistent antigen.

### Memory-like cells are enriched for common progenitors for both Tfh and Teff

To investigate the potential developmental relationships between CD4 T cell clusters, including memory-like, Teff and Tfh cells, we performed trajectory inference analysis (Monocle 3; Methods) using the scRNA-seq data. To specifically analyze the transition of cells in response to chronic antigen stimulation, we reclustered cells from c13-21 dpi and labeled the cells according to the previously defined phenotypic clusters (**Fig. 2A**). Tfh (C7r) and Teff (C8-10r) were mapped separately and most distantly in pseudotime from the naive cluster (C1r, 2r), supporting their terminally differentiated states. Naive cells were closely projected to the resting memory-like populations (C3r, 4r), but did not directly transition into either terminally differentiated population. Instead, cells in C3/4r transitioned into mature Tfh (C7r) or Teff (C8-10r) cells after passing through the activated memory-like state (C5/6r) and a subsequent bifurcation, suggesting that the memory-like state serves as a common progenitor for both Tfh and Teff fates. Along the trajectory from naive to mature Tfh cells (**Fig. 2B, top**), *Tcf7* expression remained roughly constant across all differentiation time points (**Fig. 2C**). As cells differentiated from C3-4r, *Bcl6* and *Cxcr5* expression was increased following commitment to the mature Tfh phenotype (**Fig. 2C**). In contrast, *Tcf7* expression decreased along the Teff trajectory (**Fig. 2B, bottom**), while *Runx3, Prdm1, and Ifng* expression increased (**Fig. 2D**).

**Figure 2.**
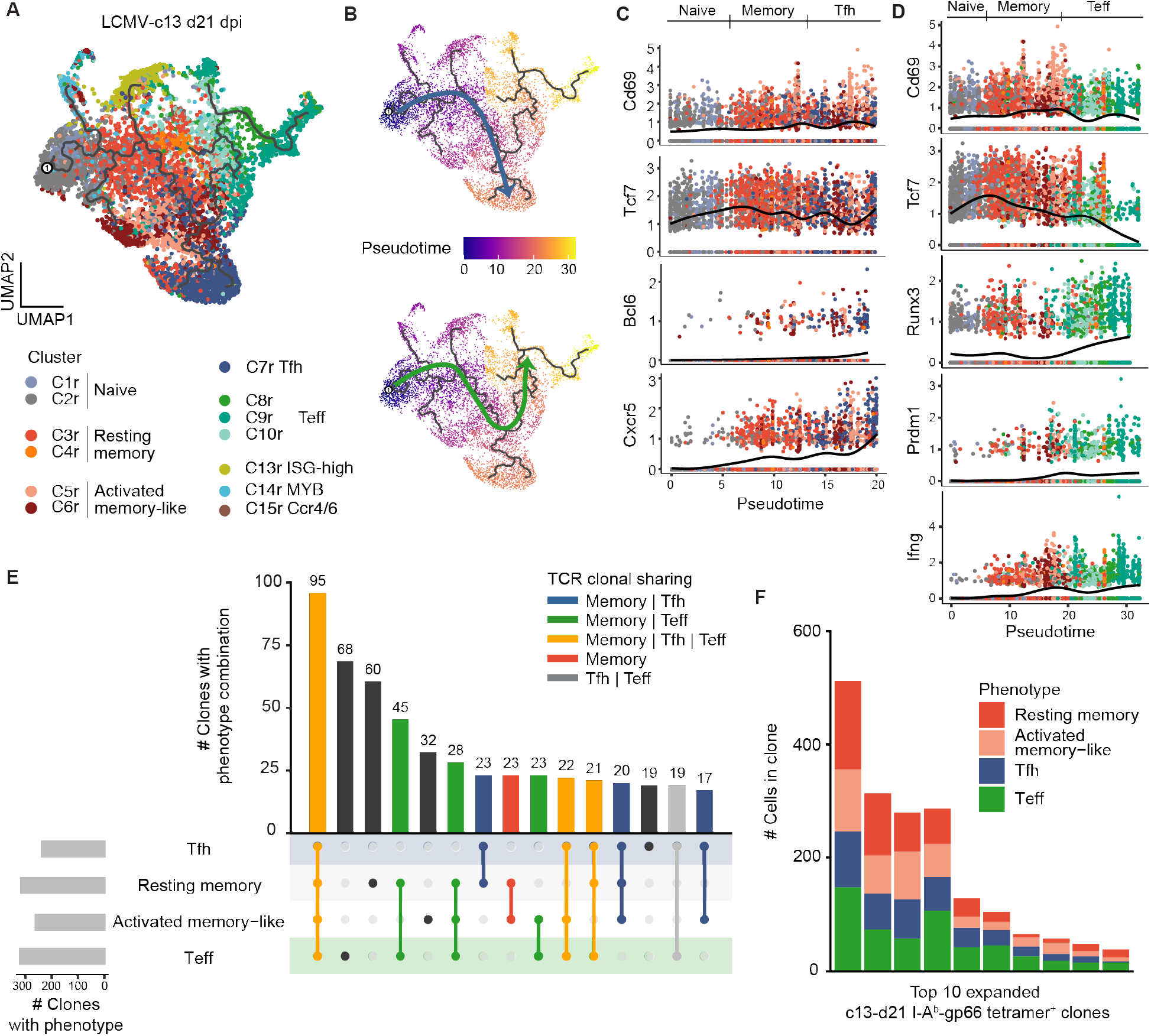
Memory-like cells are enriched for common progenitors for both Tfh and Teff. **(A)** UMAP trajectory graph of I-A^b^-gp66 tetramer-sorted LCMV-c13 21 dpi CD4 T cells excluding Tregs colored by phenotypic clusters as in Fig. 1A. **(B)** UMAP trajectory of LCMV-c13 21 dpi CD4 T cells along the Tfh branch (top) or Teff branch (bottom). Cells are colored by pseudotime. **(C)** Gene expression dynamics of activation and Tfh lineage markers along the Tfh pseudotime trajectory. **(D)** Gene expression dynamics of activation and Teff markers along the Teff pseudotime trajectory. **(E)** Phenotypic overlap of expanded (>1 cell) T cell clones among the resting memory, activated memory-like, Tfh, and Teff clusters. Bars are colored by the category of clonal phenotypic overlap. **(F)** Phenotypic composition of the top 10 most highly expanded I-A^b^-gp66 Tet^+^ clones from LCMV-c13 21 dpi.

To further support our hypothesis that cells in the C3-6r memory-like cluster constitute a common progenitor pool for Tfh and Teff cells, we paired single-cell phenotyping with TCR sequence information to define clonal developmental trajectories of memory-like (C3/4r and C5/6r), Tfh (C7r), and Teff (C8-10r) cells. We defined TCR clones based on identical TCRαβ CDR3 sequences and identified 3,837 total clones from cl13-21 dpi cells. Most of the expanded clones (>1 cell per clone) were observed in the Tet^+^ population, while the majority of Tet^−^ clones were detected as unexpanded singletons (**Fig. S2A-C**). Among expanded clones (584 total), 313 clones were found in two or more of the three large phenotype clusters and 294 of the 313 (94%) expanded clones were detected in the memory-like population, in addition to Tfh and/or Teff populations (**Fig. 2E**). Indeed, the most highly expanded clones from c13-21 dpi all comprised cells with the memory-like phenotypes (**Fig. 2F**). In contrast, only 19 expanded clones (3%) were detected in both Tfh and Teff populations, but not in the memory-like population (**Fig. 2E**). Taken together, these results suggest that during LCMV-c13 infection, the vast majority of the long-term clonal CD4 T cell response is maintained by the memory-like cells that replace the initially expanded naive pool-derived clones and function as putative common Tprog cells.

### Epigenetic differentiation trajectories of progenitor CD4 T cells

To identify epigenetic pathways of TCF-1^+^ progenitor CD4 T cell differentiation, we performed single-cell assay for transposase accessible chromatin with sequencing (scATAC-seq) on PD-1^+^ CD4 T cells from the spleens of c13-infected mice on 21 dpi. In total, we obtained high quality scATAC-seq profiles from 11,611 cells (unique fragment count >1,000, transcription start site enrichment >11; **Fig. S3A, B**) and performed dimensionality reduction using iterative latent semantic indexing followed by UMAP (Satpathy et al., 2019), which identified 8 distinct chromatin state clusters after removing thymic Tregs (**Fig. 3A**). Next, we integrated scATAC-seq and scRNA-seq datasets to define an identity of each cell cluster (**Fig. 3B; Fig. S3C**). To perform integrative analysis, we first visualized the accessibility of cell cluster marker genes identified in scRNA-seq data by calculating gene scores, which is a measurement of the overall accessibility of the gene body and surrounding open chromatin regions (OCR) (Pliner et al., 2018); see Methods). Second, we performed constrained integration of single-cell transcriptomic and chromatin accessibility profiles to calculate gene integration scores, yielding gene expression profiles at the single-cell level in the scATAC-seq defined UMAP space (see Methods). Importantly, these analyses identified cells with Tfh (C4a and C5a), progenitor (C6a and C7a) and Teff (C2a and C3a) phenotypes; C6a and 7a were marked by intermediate accessibility of *Tcf7* and *Slamf6* and low to intermediate accessibility of *Cxcr5* and *Bcl6*, corresponding to the putative Tprog cell clusters identified in the transcriptomic analysis (**Fig. 3C**). Despite the overall epigenetic similarity between C6a and C7a, hierarchical clustering of differentially activated loci revealed increased accessibility in C6a at several loci related to T cell activation, including *Cd69, Batf, Nr4a2, Irf4*, and *Tox* (**Fig. 3C; Fig. S3D, E**), suggesting that C7a and C6a correspond to the resting (C3/4r) and activated (C5/6r) Tprog cell clusters identified by scRNA-seq, respectively. To further support these findings, we defined the differential OCRs for each cluster and performed motif enrichment analysis between C6a and C7a, identifying several motifs associated with NFAT and AP-1 in C6a compared to C7a (**Fig. 3D**). In contrast, Tfh cells showed high TF motif accessibility of TCF factors, and Teff cells showed high accessibility of TBX and RUNX factors (**Fig. 3D**).

**Figure 3.**
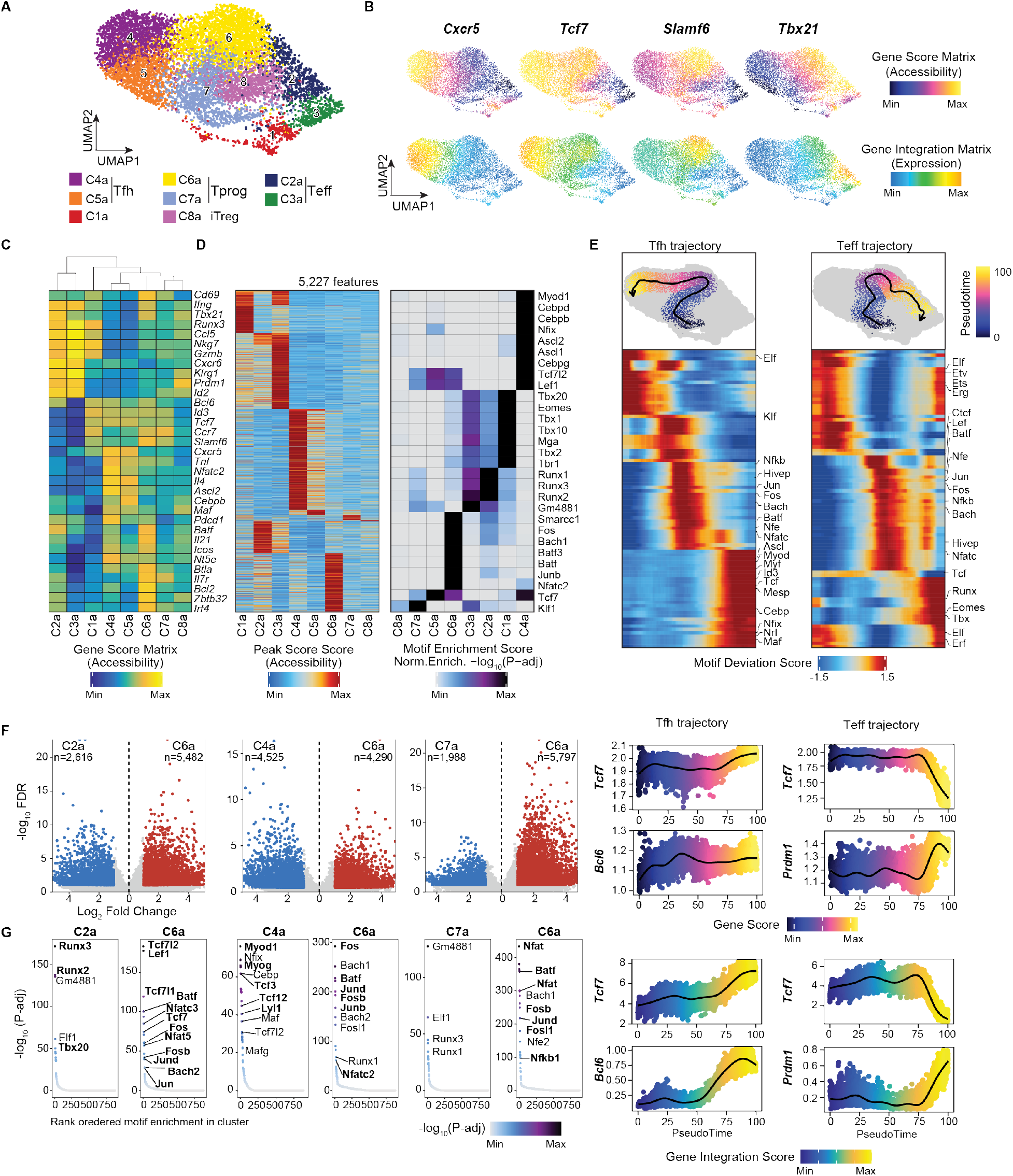
Chromatin accessibility analysis reveals heterogeneity and delineates the main CD4 differentiation pathways during chronic viral infection. **(A)** UMAP of scATAC-seq data of PD1^+^ CD4 T cells on 21 dpi of LCMV-cl13 infection. **(B)** UMAP of gene score (accessibility) and gene integration matrix (expression) for *Cxcr5, Slamf6* and *Tbx21*. **(C)** Heatmap visualization of marker gene accessibility scores across clusters represented by gene score values. **(D)** Heatmap of marker peak scores of 5,227 *cis*-regulatory elements in scATAC-seq clusters (left). Heatmap of enriched motifs in marker peaks of the specific clusters (right). **(E)** UMAP projection of Tfh and Teff differentiation trajectory, respectively. Cells that are not part of the trajectory are colored grey (top). Pseudotime heatmaps of motif deviation scores on the two differentiation trajectories (middle). Pseudotime analysis of the genescores and gene expression for indicated features (bottom). **(F)** Volcano plot visualization of the differential peak analysis between the indicated clusters (FDR <= 0.1 and a Log_2_FC >= 0.5) **(G)** Hockey plot representation of the enriched motifs under the cluster specific peak sets.

To further investigate the epigenetic dynamics between the Tprog, Teff, and Tfh populations, we constructed cellular trajectories for the transition to either terminally differentiated CD4 subset, and analyzed differential TF motif accessibility along the two paths (**Fig. 3E, top**). Since Tprog cells clustered in between the Tfh and Teff cells, and TCR-based analysis uncovered extensive clonal sharing of Teff and Tfh cells with this population, we used Tprog cells as a starting point, and reconstructed two differentiation trajectories using a nearest-neighbor approach (sequential selection of similar chromatin states based on Euclidean distance, see Methods). As cells transitioned from C7a to C6a (representing the common path of the two trajectories), they gained accessibility around NFAT, BACH, and BATF/AP-1 motifs, representing the early chromatin remodeling events of Tprog commitment. After this common differentiation step, cells bifurcated into terminally differentiated Teff and Tfh phenotypes. At these later stages of the trajectory, cells lost accessibility at common progenitor TF motifs and gained accessibility at Tfh-(e.g., TCF7, ASCL, CEBP, and MAF) (Liu et al., 2014; Shao et al., 2019; Tanaka et al., 2014; Wu et al., 2015), and Teff-related TF motifs (e.g., RUNX, TBX/EOMES) (Djuretic et al., 2007; Mullen et al., 2001; Naoe et al., 2007; Szabo et al., 2000)(**Fig. 3E**).

To quantify chromatin remodeling activities across the different phenotypic populations, we performed differential chromatin accessibility analysis between Tprog, Tfh and Th1 effector cells. Comparison of C7a and C6a Tprog cells identified 1,988 and 5,797 differentially accessible OCRs, while a comparison of C4a (Tfh) and C6a (Tprog) cells yielded 4,525 and 4,290 differential OCRs, respectively. Comparing C2a (Teff) to C6a (Tprog) cells identified 5,482 and 2,616 differentially accessible OCRs, respectively (**Fig. 3F**). We performed motif enrichment analyses at these specific OCRs and found that the Tprog populations exhibited several motifs associated with NFAT and AP-1 binding in C6a, compared to C7a, recapitulating the trajectory analysis. During the early transition from Tprog to Teff, in addition to loss of TCF7-related motifs (**Fig. 3G**) consistent with loss of Tcf7 accessibility and expression (**Fig. 3E, bottom**), NFAT and AP-1 motifs were lost. In contrast, the most significantly enriched motifs were associated with RUNX activity, which have been implicated in the commitment to the Th1 lineage (Djuretic et al., 2007; Naoe et al., 2007). During the transition from Tprog to Tfh, we detected a similar reduction of accessibility to NFAT and AP-1 motifs, while several motifs associated with the binding of basic helix-loop-helix (bHLH) proteins were enriched, which have been shown to promote Tfh differentiation at the expense of Th1/Teff differentiation (Liu et al., 2014; Shaw et al., 2016).

Altogether, these results suggest that activated CD4 T cells adopt two distinct states of TCF-1^+^ memory-like phenotypes: resting Tcmp-like (C3/4r and C7a) and activated memory (C6a) states. Resting memory-like cells are mobilized to terminal differentiation by transient TCR signaling initially without polarization towards either lineage, and the specification of each terminal cell type is likely initiated after cells complete the transition to C6a through the activation of lineage-specific TF binding motifs. These two-step processes may be distinct from the initial differentiation towards either lineage as an immediate consequence of T cell activation during the initial priming of naive T cells.

### TCF-1^+^ BCL6^lo/−^ cells exhibit superior proliferative capacity and give rise to both TCF-1^−^ Teff and TCF-1^+^ BCL6^hi^ Tfh following adoptive transfer

To demonstrate the progenitor function of the TCF-1^+^ BCL6^lo/−^ cells *in vivo*, we compared repopulation and differentiation capacities of the three major subsets of PD-1^+^ CD4 T cells isolated from LCMV-c13 infected mice. Each population was adoptively transferred into congenic B6-CD45.1 mice, which were either subsequently infected with LCMV-c13, or infection-matched at the time of transfer (**Fig. 4**). We used Ly-108 and CXCR5 as surrogate surface markers for the expression of TCF-1 and BCL6 (**Fig. S1E**). Ly-108^+^ CXCR5^−^ (enriched for TCF-1^+^ BCL6^lo/−^ cells), Ly-108^+^ CXCR5^+^ (enriched for TCF-1^+^ BCL6^hi^ cells), and Ly-108^−^ CXCR5^−^ (enriched for TCF-1^−^ BCL6^−^ cells) PD-1^+^ CD4 T cells were harvested as donor cells from B6 mice that had been infected with LCMV-c13 for 21-24 days.

**Figure 4.**
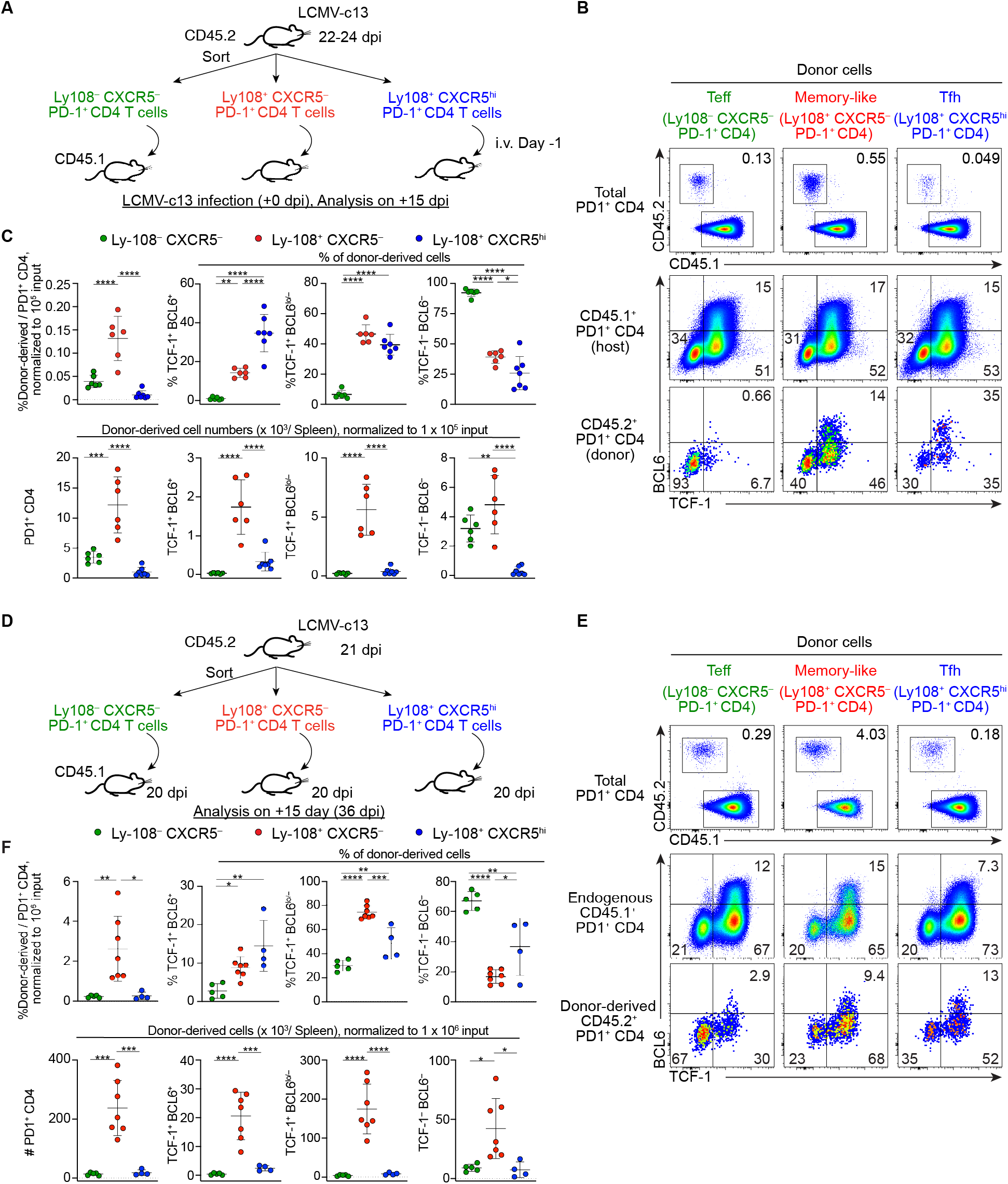
TCF-1^+^ BCL6^lo/−^ PD-1^+^ CD4 T cells are capable of superior expansion and differentiation into both Teff and Tfh following adoptive transfer. **(A-C)** (A) shows the experimental design of adoptive transfer of distinct populations of PD-1^+^ CD4 T cells from LCMV-c13 infected mice to naive mice subjected to subsequent infection with LCMV-c13. Representative flow cytometry data showing frequencies of donor-derived cells and their expression of TCF-1 and BCL6 15 days after infecting the recipient mice with LCMV-c13 were shown in (B). Pooled data from 2 experiments with n = 3 / transfer group / experiment are shown in (C) with mean±SD with statistical analysis with One-way ANOVA. **(D-F)** Experimental design of adoptive transfer of distinct populations of PD-1^+^ CD4 T cells from LCMV-c13 infected mice to infection-matched recipient mice (A). Representative data and statistical data from 2 experiments with n = 2-4 / transfer group / experiment are shown in B-C.

Following transfer to naive host mice and a subsequent challenge with LCMV-c13 infection (**Fig. 4A**), the donor-derived Ly-108^+^ CXCR5^−^ cells expanded at a significantly higher magnitude compared to the two other populations of donor cells (**Fig. 4B, C**). They repopulated all three major populations of the PD-1^+^ CD4 T cell pool at similar ratios compared to those of endogenous CD4 T cells (**Fig. 4B, middle, and C**), indicating their progenitor function. In contrast, Ly-108^−^ CXCR5^−^ Teff cells remained predominantly as TCF-1^−^ BCL6^−^ cells and generated few TCF-1^+^ CD4 T cells with low overall proliferation (**Fig. 4B, left, and C**), an indication of their terminal differentiation. Ly-108^+^ CXCR5^hi^ Tfh cells expanded the least among the three donor populations and gave rise to a higher frequency of TCF-1^+^ BCL6^+^ cells compared to the other populations (**Fig. 4B, right, and C**). We detected TCF-1^+^ BCL6^−^ and TCF-1^−^CD4 T cells in the mice that had received donor Tfh cells; however, the absolute numbers of these cells were much lower than those derived from donor Teff or memory-like cells (**Fig. 4C**). Therefore, it is likely that these non-Tfh cells were derived from low frequencies of contaminating Teff or memory-like cells in the Tfh donor cells.

To more stringently test the ability of cells in each CD4 subset to proliferate and differentiate, we performed transfers to infection-matched recipient mice (**Fig. 4D-F**). Spleens from recipient mice were analyzed 15 days after transfer (36 dpi), at which time viremia still remained high (data not shown). Similar to the transfer to naive host mice followed by rechallenge, Ly-108^+^ CXCR5^−^ cells demonstrated the highest proliferation capacity with more than 10-fold expansion compared to Teff and Tfh donor cells (**Fig. 4E, F**). They gave rise to all three populations at a ratio similar to endogenous CD4 T cells, further demonstrating their lineage plasticity and self-maintenance. In contrast, although both transferred Teff and Tfh cells gave rise to cells distinct from the pre-transferred phenotypes, they both expanded at substantially lower magnitudes compared to memory-like donor cells. Thus, it is likely that the observed phenotypic changes were caused by infrequent, contaminating memory-like cells during purification (**Fig. 4E, F**).

Together, the results from these adoptive transfer experiments demonstrated that the Ly-108^+^ CXCR5^−^ activated CD4 T cells retain high proliferative capacity and plasticity to differentiate into both TCF-1^−^ Teff and TCF-1^+^ Tfh cells. These results suggest that they act as progenitor cells to maintain the pool of differentiated Teff and Tfh cells that are terminally differentiated and exhibit limited proliferative potential.

### *TCF-1*^+^ BCL6^lo/−^ *PD-1^+^ CD4 cells are progenitors that maintainTCF-1^−^ effector cells in vivo*

Our results thus far have shown that in the presence of chronic antigen, CD4 T cells preferentially adopt the TCF-1^+^ BCL6^lo/−^ phenotype after a wave of binary differentiation from naive CD4 T cells into Teff and Tfh cells. TCF-1^+^ BCL6^lo/−^ cells can give rise to TCF-1^−^ Teff cells following adoptive transfer and thus can function as progenitors for antigen-specific CD4 T cells. To determine whether they are required for sustained CD4 Teff response, we conditionally deleted *Bcl6* in a CD4 T cell-specific manner. To achieve this, we generated a novel *cre* transgenic driver that facilitates deletion of a *loxP*-flanked gene in CD4 T cells - with minimal impact on CD8 T cells - by knocking-in *cre* into the *Cd40lg* locus on the X chromosome (**Fig. S4A**). This cre driver deleted a *loxP*-flanked transcriptional stop cassette in the *Rosa26* locus in virtually all CD4^+^ T cells, whereas its activity in CD8 T cells was substantially lower and absent in B cells, myeloid cells, and NK cells (**Fig. S4B-F**). Since germinal center B cells were formed comparably in *CD40lg-cre* male mice compared to control mice, *cre* knock-in to the 3’ UTR preserved the function of the *Cd40lg* gene (**Fig. S4G**).

Using this animal model, we investigated the role of *Bcl6* in the maintenance of long-term antigen specific CD4 T cell response. Interestingly, 8 days following LCMV-c13 infection, the number of Tet^+^ cells was comparable in *Cd40lg-cre Bcl6*^F/F^ compared to control *Cd40lg-cre Bcl6^+/+^* mice (**Fig. 5A, B**). In Tet^+^ CD4 T cells, TCF-1^+^ cells were practically absent in *Cd40lg-cre Bcl6*^F/F^ mice with the number of TCF-1^−^ CXCR6^+^ Teff cells remaining intact (**Fig. 5C, D**). This result indicates that the deletion of *Bcl6* was complete and that the development of both TCF-1^+^ BCL6^hi^ Tfh and TCF-1^+^ BCL6^lo/−^ cells requires *Bcl6*, confirming the the requirement for *Bcl6* in memory CD4 T cells and Tfh cells (Choi et al., 2013; Ichii et al., 2007; Pepper et al., 2011). We also observed a similar phenotype in LCMV-Arm infected mice, where *Bcl6* deletion caused not only a loss of Tfh cells as defined by CXCR5^+^ PD1^+^ CD4 T cells or I-A^b^-gp66 Tet^+^ TCF-1^+^ BCL6^hi^ cells, but also a loss of TCF-1^+^ BCL6^lo/−^ cells, which were enriched for CCR7^+^ Tcmp cells (**Fig. S5A, B**). The CD4 Teff cells in LCMV-c13-infected *Cd40lg-cre Bcl6*^F/F^ mice and control mice expressed comparable levels of the inhibitory receptor TIM3, indicating they are qualitatively similar (**Fig. S5C, D**). Although the number of Teff cells was comparable between *Cd40lg-cre Bcl6*^F/F^ and control mice on 8 dpi, Tet^+^ cells were reduced more than 10-fold by 15 dpi in *Cd40lg-cre Bcl6*^F/F^ mice (**Fig. 5C, D**), indicating that the initially generated BCL6-independent Teff cells derived from naive CD4 T cells are short-lived and that *Bcl6* and BCL6-dependent TCF-1^+^ CD4 T cells are essential for the maintenance of BCL6^−^ TCF-1^−^ Teff cells.

**Figure 5.**
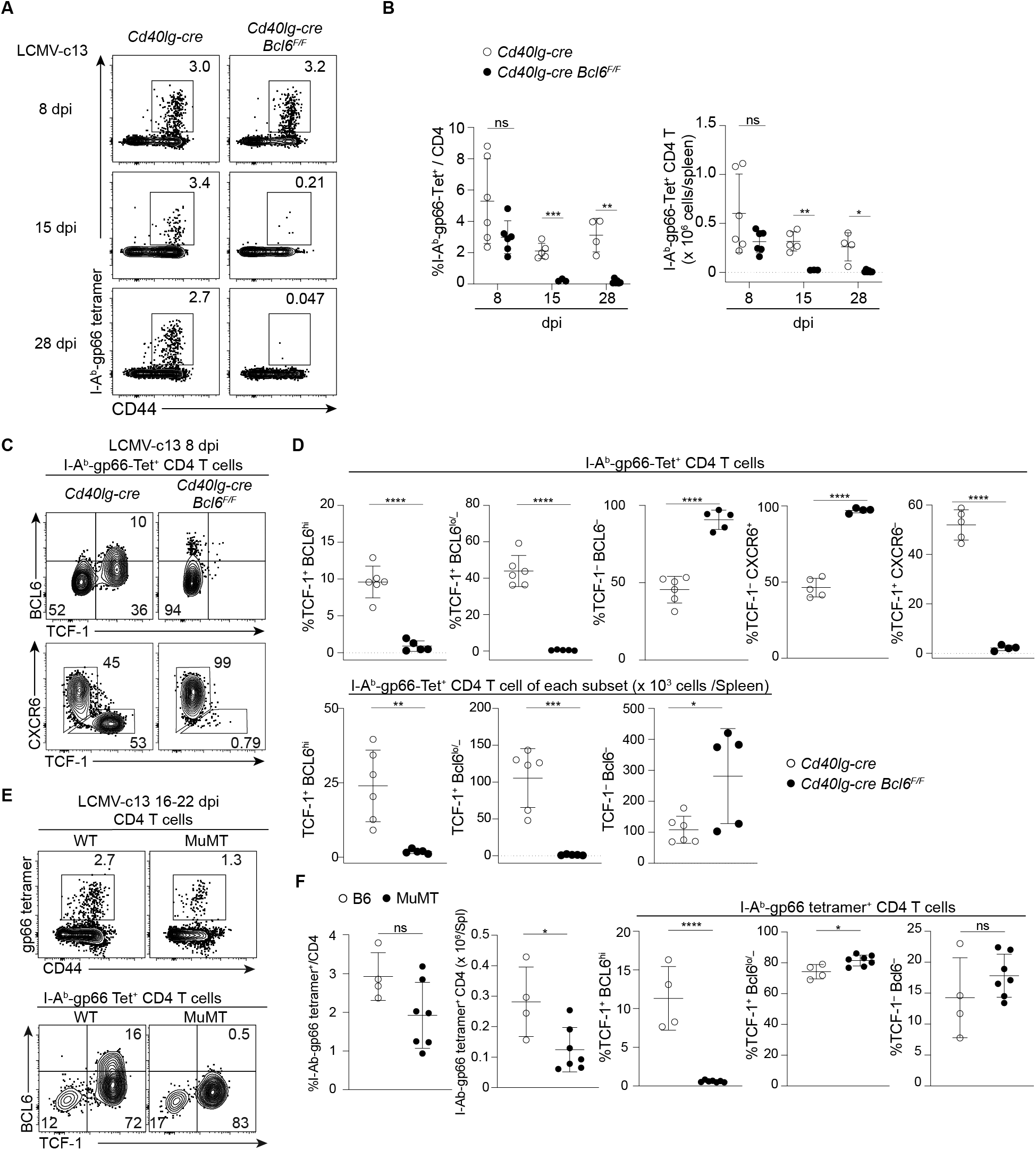
The maintenance of TCF-1^−^ Teff cells requires BCL6-dependent, B cell-independent, TCF-1^−^ BCL6^lo/−^ PD-1^+^ memory-like CD4 T cells. **(A-B)** Representative flow cytometry plots showing expression of CD44 and I-A^b^-LCMV-gp66-specific TCR in splenic CD4 T cells in *Cd40lg-cre Bcl6*^F/F^ and control *Cd40lg-cre* mice infected with LCMV-c13, analyzed at 8 dpi, 15 dpi or 28 dpi. Pooled data from 2 experiments with 2-3 mice/group/time point are shown with mean±SD. Unpaired *t*-test. **(C-D)** Expression of TCF-1, BCL6 and CXCR6 in I-A^b^-gp66-specific splenic CD4 T cells from LCMV-c13-infected with *Cd40lg-cre Bcl6*^F/F^ and control *Cd40lg-cre* mice on 8 dpi. Data pooled from two experiments with n = 2-4 per genotype are shown with mean±SD. **(E-F)** Expression of TCF-1, BCL6 and CXCR6 in I-A^b^-gp66-specific splenic CD4 T cells from LCMV-c13-infected with MuMT and age-matched B6 (WT) mice on 16-22 dpi. Data from two experiments with n = 2-5 per genotype per experiment are shown with mean±SD.

The TF Blimp1 plays antagonistic roles to BCL6 in the differentiation of Tfh versus Teff CD4 T cells, memory versus effector CD8 T cells, and memory B versus plasma cells (Kallies et al., 2009; Nutt et al., 2007; Rutishauser et al., 2009; Shaffer et al., 2002; Shao et al., 2019; Shapiro-Shelef et al., 2003; Welsh, 2009; Wu et al., 2015). In the absence of Blimp1 in *Cd40lg-cre Prdm1*^F/F^ mice, expansion of Tet^+^ CD4 T cells was increased by 3-fold compared to control mice on 8 dpi with LCMV-c13 infection (**Fig. S5E, F**). In addition to the greater expansion, the vast majority of Tet^+^ CD4 T cells in *Cd40lg-cre Prdm1*^F/F^ mice were TCF-1^+^. Thus, Blimp1 not only regulates the expansion of antigen-specific CD4 T cells, but also promotes the differentiation of TCF-1^+^ into TCF-1^−^ CD4 T cells.

To determine whether TCF-1^+^ BCL6^hi^ Tfh cells are necessary for the development of TCF-1^+^ BCL6^lo/−^ cells or the maintenance of TCF-1^−^ Teff cells, we analyzed MuMT mice lacking TCF-1^+^ BCL6^hi^ cells due to B cell deficiency. Total Tet^+^ cells were mildly diminished on 16 dpi or later and the frequency of cells in the TCF-1^+^ BCL6^lo/−^ compartment was comparable to control WT mice despite the total loss of TCF-1^+^ BCL6^hi^ cells (**Fig. 5E, F**). Together with the limited expansion capacity in adoptive transfer experiments shown in Fig. 4, these results suggest that BCL6^hi^ CXCR5^hi^ cells are dispensable for the maintenance of TCF-1^+^ BCL6^lo/−^ and TCF-1^−^ PD-1^+^ CD4 T cells in response to chronic LCMV-c13 infection. Our results establish *Bcl6* as a critical regulator of the differentiation of TCF-1^+^ BCL6^lo/−^ Tprog cells, which are indispensable for long-term antigen specific CD4 T cell response during chronic viral infection. Additionally, these findings raise the question whether a similar progenitor population might support sustained antigen specific CD4 responses during anti-tumor immune responses.

### Tumor antigen-specific CD4 T cells differentiate predominantly into TCF-1^+^ BCL6^lo/−^ cells in tumor-draining lymph nodes

In order to examine CD4 T cell differentiation in anti-tumor immunity and determine if it shows similarities to chronic LCMV infection, we examined the differentiation and proliferation of OT-II CD4 T cells expressing an I-A^b^-restricted OVA-specific TCR in response to the subcutaneously transplanted ovalbumin (OVA) expressing 1956 sarcoma cell line (Ferris et al., 2020). We adoptively transferred CFSE-labeled, CD45.1/2 OT-II CD4 T cells one day prior to tumor inoculation and tracked their proliferation by CFSE dilution and differentiation by staining for TCF-1 and BCL6 in the tumor and tdLNs (**Fig. 6A**).

**Figure 6.**
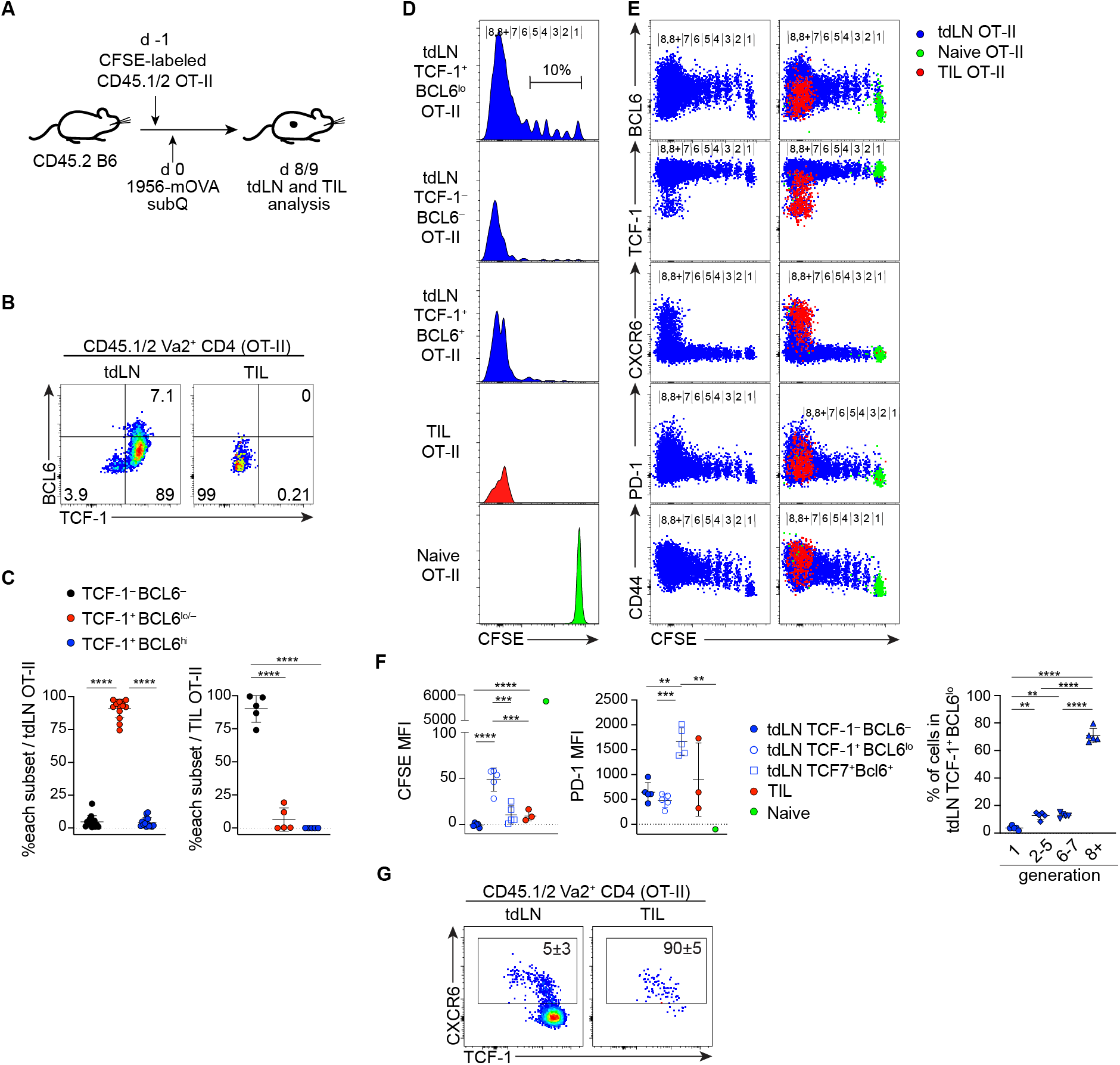
Antigen-specific CD4 T cells differentiate into TCF-1^+^ PD-1^+^ cells following extensive division in tumor-draining lymph nodes, but not in the tumor microenvironments. **(A)** Experiment design to examine CD4 T cell response to tumor antigen. **(B-C)** Representative flow plots showing expression of TCF-1 and BCL6 by donor-derived OT-II CD4 T cells harvested from tumor (TIL) and the tumor draining lymph node (tdLN). Data are pooled from two experiments (n = 6 - 8 per experiment) and shown with mean±SD. **(D-F)** CFSE dilution and expression of BCL6, TCF-1, CXCR6, PD-1 and CD44 of donor-derived OT-II cells. The CFSE level of naive OT-II cells was determined by control recipient mice without tumor transplantation sacrificed at the same time points. Left panels in (E) show only tdLN-derived OT-II cells without overlay of TIL or naive cells. Pooled data are shown with mean±SD in (F) with assessment of statistical differences by one-way ANOVA. **(G)** Representative flow cytometry plots showing expression of CXCR6, TCF-1 and BCL6 by donor-derived OT-II CD4 T cells harvested from tdLN and the tumor. Numbers shown are mean±SD.

In tdLNs, OT-II CD4 T cells differentiated into all three populations defined by the expression of TCF-1 and BCL6 (**Fig. 6B, C**). As we observed in LCMV-c13 infected mice, the frequency of the TCF-1^+^ BCL6^lo/−^ cells was the highest among expanded OT-II cells. Although TCF-1^−^ Teff, BCL6^+^ Tfh and the majority of TCF-1^+^ BCL6^lo/−^ cells fully diluted CFSE over several days, approximately 10% of TCF-1^+^ BCL6^lo/−^ cells underwent limited rounds of division and expressed higher levels of BCL6 compared to fully divided TCF-1^+^ cells (**Fig. 6D-F**). Thus, the tumor-reacting TCF-1^+^ BCL6^lo/−^ cells were heterogeneous, ranging from BCL6^int/hi^ PD-1^lo^ dormant cells to extensively divided BCL6^lo/−^ PD-1^+^ cells, which resembled Tprog found in LCMV-infected mice. In contrast to cells in tdLNs, OT-II cells isolated from the tumors were predominantly TCF-1^−^ CXCR6^+^ Teff cells (**Fig. 6B, C**). Since a small fraction of extensively divided OT-II CD4 T cells in tdLN upregulated CXCR6 before downregulating TCF-1 expression **(Fig. 6G)**, tumor-reactive CD4 T cells likely initiate their Teff differentiation during the TCF-1^+^ state prior to migration to the tumor microenvironment. These results suggest that in the setting of chronic antigen stimulation, rather than binary differentiation to TCF-1^−^ BCL6^−^ or TCF-1^+^ BCL6^+^ populations, CD4 T cells preferentially differentiate into TCF-1^+^ BCL6^lo/−^ cells, which may serve as progenitors and continuously generate fully differentiated Teff and Tfh cells as they continue to recognize antigen. These results also suggest that such continued differentiation mainly takes place in tdLNs rather than in tumors, possibly because only the tdLNs contain the appropriate microenvironment for TCF-1^+^ BCL6^lo/−^ cells to retain a partially differentiated phenotype.

## Discussion

Using a combination of single-cell genomics and population-based experiments in the chronic LCMV-c13 model, we demonstrate that antigen-specific CD4 T cells differentiate into TCF-1^+^ BCL6^lo/−^ PD-1^+^ CD4 T cells that serve as progenitor cells and give rise to Teff and Tfh cells in response to persistent antigen. During an acute infection or in response to vaccination, significant proportions of Teff and Tfh cells are derived directly from naive CD4 T cells (Choi et al., 2011; DiToro et al., 2018). In contrast, CD4 T cell response to chronic antigen is more complicated and requires progenitor cells that can support long-term Teff responses as the initial Teff cells decay. This process is illustrated in our *Cd40lg-cre Bcl6*^F/F^ model, where the initial Teff cell expansion from naive cells was intact, but Teff cells fail to persist due to a lack of Tprog cells that generate new Teff cells. Thus, the developmental pathway of CD4 Teff cells is shifted from BCL6-independent to -dependent as antigen persists.

The progenitor CD4 T cells identified in this study exhibit similarities to CD4 Tcm, Tcmp, and pre-Tfh cells that also express *Tcf7, Slamf6*, and *Bcl2* (Choi et al., 2011; Ciucci et al., 2019; Pepper and Jenkins, 2011). Shortly after activation, ranging from several hours to a few days, CD4 T cells diverge into IL2Ra^+^ Blimp1^+^ Teff cells and IL2Ra^−^ BCL6^+^ cells in a B-cell independent manner (Baumjohann et al., 2011; Choi et al., 2011). These IL2Ra^−^ BCL6^+^ cells contain Tcmp and pre-Tfh (Choi et al., 2013). This binary fate choice between Teff and Tfh is stable, since differentiation of adoptively transferred IL2Ra^−^ BCL6^+^cells is substantially biased towards Tfh, and the depletion of these early IL2Ra^−^ BCL6^+^ cells causes a significant reduction of Tfh cells (Choi et al., 2013; DiToro et al., 2018). In the contexts of acute infection or immunization with non-replicating antigen, both antigen levels and inflammation wane by the time pre-Tfh or Tcmp cells develop. In contrast, following LCMV-c13 infection, antigen levels remain high for the first several days, rendering TCF-1^+^ PD-1^+^ CD4 T cells that might otherwise become pre-Tfh or Tcmp exposed to continued signals through TCR and other receptors. Such continued stimulation may result not only in changes in surface marker expression, such as increased PD-1 and reduced CCR7 and CXCR5 compared to Tcmp (Ciucci et al., 2019), but also epigenetic changes that preserve Teff differentiation.

Our single-cell analysis revealed two states of cells in the progenitor pool with distinct activation signatures. The transition between the two states is associated with NFAT- and AP-1-target gene activation; increased chromatin accessibility of these TF binding motifs was the dominant change between the two states. Since Tfh differentiation is more pronounced than Teff differentiation in CD4 T cell responses to chronic LCMV infection, it is possible that the progenitor CD4 T cells in the resting state have already been biased towards Tfh or they robustly acquire a Tfh-biased gene expression signature as they are mobilized to the activated states. However, trajectories between the resting progenitor state to Tfh or Teff using both transcriptomic and epigenetic analyses suggest that the specification to either lineage occurs independently of the initial transition from the resting to activated states. Activation of E-protein targets, which may be mediated by E2A and ASCL2 (Liu et al., 2014; Shaw et al., 2016), in progenitor-Tfh tranjectory and activation of RUNX targets for the Teff trajectory (Djuretic et al., 2007; Naoe et al., 2007) were detected as progenitor CD4 T cells acquire transient NFAT and AP-1 activation. Although a stringent validation may require fate tracing at the single cell level, this conclusion is also supported by the substantially overlapping TCR clonality among resting and activated Tprog, Tfh and Teff populations. The activated Tprog cells may overlap with previously described, BCL6-dependent PD-1^+^ CD4 T cells that develop during *M. tuberculosis* infection (Moguche et al., 2015).

Finally, our study highlights the similarities between progenitor CD4 and CD8 TPEX cells. Both cell types develop in the presence of persistent antigen, express TCF-1, Ly-108, and PD-1, and continue generating differentiated effectors while they self-renew. While these cell types resemble CD4 Tcmp and CD8 MPEC, respectively, they have unique gene expression and epigenetic signatures, such as those associated with activation. In addition, we observed the emergence of PD-1^+^ CD4 T cells in tumor-bearing mice, which phenocopy progenitor CD4 T cells in response to LCMV infection. Many studies have demonstrated the importance of CD8 TPEX in enabling durable anti-tumor immunity and the immune response to checkpoint blockade (ICB) therapies. CD8 TPEX are found in tumors and extratumoral tissues, such as lymph nodes. In mouse tumor transplantation models, it has been demonstrated that intratumoral CD8 TPEX cells are sufficient to promote anti-tumor immunity in response to vaccine or ICB (Siddiqui et al., 2019). In our tumor transplantation experiments, we found tumor-reactive TCF-1^+^ PD-1^+^ CD4 T cells exclusively in the tdLNs while almost all antigen-specific CD4 T cells in the tumor were differentiated TCF-1^−^ Teff cells. In the lymph nodes, a small fraction of tumor-reactive TCF-1^+^ CD4 T cells initiate Teff differentiation as indicated by CXCR6 upregulation. The roles of CD4 T cells to ICB in direct antitumor immunity or in support of CD8 T cell function invoked by ICB is not clearly understood. However, the mobilization of CD8 TPEX by ICB in tumor-draining lymph nodes could be more efficacious for certain types of tumors which require CD4 T cells for rejection through co-activation of both CD4 and CD8 progenitor populations.

In summary, our analysis decoded the heterogeneity of CD4 T cells that respond to persistent antigen in the context of antiviral and anti-tumor immunity and highlighted a population of TCF-1^+^ BCL6^lo/−^ PD-1^+^ CD4 T cells as progenitor cells that support the continued generation of differentiated effectors and helper CD4 T cells. Progenitor CD4 T cells go through a transitory state in which they retain an unbiased epigenetic signature towards either terminal fate, and subsequently resolve the bipotential states, which is distinct from the binary differentiation of naive CD4 T cells into Teff and Tfh early in an immune response. Our results reveal the population dynamics and differentiation hierarchy of CD4 T cells for their sustained responses.

## Acknowledgments

We thank the NIH tetramer core at Emory for providing the I-A^b^-LCMV-gp66 tetramers. This study was supported by NIH grants R01AI130152 (to T.E.), R21AI161040 (to T.E.), 5T32AI007290 (to J.A.P.), K08CA230188 (to A.T.S.), U01CA260852 (to A.T.S.), R01CA190700 (to R.D.S), P30AR073752 (Rheumatic Diseases Research Resource-Based Center at Washington University), the Leukemia and Lymphoma Society Scholar Award (to T.E.), the Parker Institute for Cancer Immunotherapy (to A.T.S. and R.D.S), the Burroughs Wellcome Fund Career Award for Medical Scientists (to A.T.S.), a Technology Impact Award from the Cancer Research Institute (to A.T.S.), and a Pew-Stewart Scholars for Cancer Research Award (to A.T.S.).

## Author Contributions

YX, KS, AS and TE designed the study. YX, KS, YQ, SR, RW and TE conducted experiments with support from BD, YQ, YT, HS and MO for data collection. KS and JP analyzed the single cell data. YX, KS, JP, BD, AS and TE interpreted results. RDS and KMM provided experimental material. YX, KS, JP, BD, AS and TE wrote the manuscript with comments from all authors.

## MATERIAL and METHODS

### Mice and infection

Male C57BL/6N and B6-CD45.1 mice were purchased from Charles River Laboratories and JAX. *Prdm1-EYFP, Bcl6*-flox mice were obtained from the Jackson Laboratory. *Cd40lg-cre* mice were generated by knocking in a mammalian codon optimized cre coding sequence following an internal ribosomal entry sequence (IRES) into 3’ UTR of the Cd40*lg* locus by homologous recombination in JM8N4 embryonic stem cells. After germline transmission, the FRT-flanked selection cassette was removed by crossing to *Actb-Flpe* transgenic mice (JAX). Because the *Cd40lg* locus is on X-chromosome, all female mice used in the *Bcl6* deletion experiments were homozygous for cre knock-in. All mice were housed in a specific pathogen-free facility at Washington University in St. Louis and were used for infection at 8–12 wk of age, unless stated otherwise. LCMV infection was performed essentially as described (Chou et al., 2016). All experiments were performed according to a protocol approved by Washington University’s Institutional Animal Care and Use Committee.

### scRNA-seq and TCR-seq sample and library generation

Single-cell RNA-seq libraries were prepared using the 10X Chromium Next Gem Single Cell V(D)J Reagent Kit (v1.1 Chemistry), according to the manufacturer’s instructions. Briefly, FACS sorted cells were washed once with PBS + 0.04% BSA and resuspended in PBS containing 0.04% BSA. Following reverse transcription and cell barcoding in droplets, emulsions were broken, and cDNA purified using Dynabeads MyOne SILANE followed by PCR amplification (98°C for 45 sec; 14 cycles of 98°C for 20 sec, 67°C for 30 sec, 72°C for 1 min; 72°C for 1 min). Amplified cDNA was then used for both 5′ gene expression library construction and TCR enrichment. For gene expression library construction, 50 ng of amplified cDNA was used for fragmentation, following by and end-repair, double-sided size selection with SPRIselect beads, PCR amplification with sample indexing primers (98°C for 45 sec; 14 cycles of 98°C for 20 sec, 54°C for 30 sec, 72°C for 20 sec; 72°C for 1 min), and double-sided size selection with SPRIselect beads. For TCR library construction, TCR transcripts were enriched from 2 μl of amplified cDNA by PCR (primer sets 1 and 2: 98 °C for 45 s; 10 cycles of 98 °C for 20 s, 67 °C for 30 s, 72 °C for 1 min; 72 °C for 1 min). Following TCR enrichment, 5 - 50 ng of enriched PCR product was fragmented and end-repaired, size-selected with SPRIselect beads, PCR-amplified with sample-indexing primers (98 °C for 45 s; 9 cycles of 98 °C for 20 s, 54 °C for 30 s, 72 °C for 20 s; 72 °C for 1 min), and size-selected with SPRIselect beads._scRNA/TCR-seq libraries were quantified using a Qubit dsDNA HS Assay kit (Invitrogen) and a HighSensitivity DNA chip run on a Bioanalyzer 2100 system (Agilent). Sequencing was performed on NovaSeq S4 (Illumina) with paired-end reads (2 x 150 cycles).

### scRNA-seq and TCR-seq library processing

Reads from 10x scRNA expression libraries were aligned to mouse genome assembly GRCm38 (mm10) and quantified using cellranger *count* (10x Genomics, version 3.1.0). The filtered feature-barcode matrices containing only cellular barcodes were used for further analysis. Single cell gene expression matrices were imported into R (version 3.6.1) and analyzed using Seurat (version 3.1.1) (Stuart et al., 2019). Cells for which the number of genes captured fell within two standard deviations of the mean were kept. Additionally, cells with greater than 5% mitochondrial RNA reads were excluded from subsequent analyses.

Single cell TCR reads were aligned to mouse genome assembly GRCm38 (mm10) and assembled into reconstructed TCR consensus sequences using cellranger *vdj* (10x Genomics, version 3.1.0). Only productive TCRα and TCRβ sequences were considered for further analysis. Overall, TCR sequences were annotated for 20,998 cells that passed RNA quality filtering, with paired TCRαβ sequences detected for 20,092 cells (95.7%). Only cells with conventional paired TCR chain combinations αβ or ααβ were kept for downstream analyses. Cells sharing the same CDR3αβ amino acid sequences were defined as belonging to the same TCR clone.

### scRNA-seq data integration and clustering

scRNA-seq libraries of I-A^b^-LCMV-gp66 Tet^+^-sorted CD4 T cell populations from Arm-8 dpi and cl13-8 dpi and −21 dpi were normalized individually using SCTransform while regressing out percent mitochondrial reads. Samples were then integrated by identifying anchors between datasets using 30 CCA dimensions. TCR genes were excluded from the selection of integration anchors to prevent TCR chain driven biases. Dimensionality reduction of the integrated matrix was performed using Uniform Manifold Approximation and Projection (UMAP) with the first 30 principal components. Phenotypic clusters were defined by constructing a k-nearest neighbors graph and identifying groups of cells using the Louvain algorithm with resolution of 0.6.

### Trajectory inference

To perform trajectory analysis of the CD4 T cell response at c13-21 dpi, dimensionality reduction of I-A^b^-LCMV-gp66 Tet^+^ and Tet^−^ scRNA-seq samples was performed using UMAP as described above after excluding cells belonging to the Treg clusters C11/12r. Pseudotime analysis was then performed with Monocle 3 (Cao et al., 2019) by learning a principal graph for the data and ordering cells along the graph using the cells in the naive phenotype cluster to select a root node.

### scATAC-seq sample and library generation

Single cell ATAC-seq experiments were performed on the 10x Chromium platform as described previously (Satpathy et al., 2019). Briefly, following sorting, cells were subjected to nuclei isolation according to the manufacturer’s recommendation. After tagmentation, nuclei were processed for generating scATAC-seq libraries and loaded to the 10x Chromium controller. For GEM incubation the standard thermocycler conditions were used and library construction was done as described by 10x Genomics for scATAC-seq. Libraries were quantified using a Qubit dsDNA HS Assay kit (Invitrogen) and a HighSensitivity DNA chip run on a Bioanalyzer 2100 system (Agilent). Sequencing was performed on NovaSeq S4 (Illumina) with paired-end reads (2 x 150 cycles), and demultiplexed using CellRanger-ATAC v1.2.

### scATAC-seq analysis

scATAC-seq datasets were processed as described previously (Granja et al., 2021). Briefly, reads were filtered, trimmed, and aligned to the mm10 reference genome using the 10x cellranger atac-count pipeline. Fragment files were loaded into ArchR for additional processing and analysis. Doublets were identified and removed using ArchR’s default doublet simulation and calling procedures. Barcodes were removed that had an enrichment of Tn5 insertions in transcription start sites (TSS enrichment) less than 4 or less than 1000 fragments. Tiles and GeneScores matrices were computed by summing Tn5 insertions in predefined genomic windows. After clustering the cells, peaks were called by macs2 (Zhang et al., 2008) on pseudoreplicates sampled from each cluster to obtain a reproducible peak set retaining cell type specific peaks. TF motif deviations were computed using chromVar (Schep et al., 2017). Imputation was performed using Magic (van Dijk et al., 2018).

### Cell preparation, cell staining, and flow cytometry

Single-cell suspensions of splenocytes were prepared by manual disruption with frosted glass slides. Bone marrow cells were dissociated from the femur with mortar and pestle. Liver cells were prepared by manual disruption with frosted glass slides followed by gradient centrifugation with 40% Percoll. For labeling of circulating cells, 3 ug biotinylated CD3e were injected i.v. and the mouse was euthanized after 3 minutes and tissue harvested.

Absolute live cell counts were determined by trypan blue exclusion using Vi-CELL (Beckman Coulter). PE- or APC-labelled I-A^b^-LCMV-gp66-77 tetramer reagents were obtained from the NIH Tetramer Core at Emory and cells were stained at room temperature for 60 min. Staining for TCF-1, BCL6, and Ki67 was performed using the Foxp3 staining kit (eBioscience) according to the manufacturer’s instructions. The following antibodies were purchased from BioLegend, unless otherwise indicated:

Alexa Fluor (AF)-conjugated donkey polyclonal anti-rabbit IgG (Thermo Fisher Scientific, catalog no. R37118); AF647–conjugated goat polyclonal anti-rabbit IgG (Cell Signaling, catalog no. 4414S); AF700-conjugated anti-CD44 (IM7), anti-CD45.2 (A20); APC-conjugated anti-PD-1 (29F.1A12), anti-CD366 (TIM-3)(RMT3-23); APC-Cy7-conjugated anti-CD8a (53-6.7, BD Biosciences), BV421-conjugated anti-CXCR5 (L138D7); Biotinylated anti-CD3e (145-2C11); PE-conjugated anti-GL7 (GL7); PerCP-Cy5.5-conjugated anti-CD4 (GK1.5), anti-CD45.1 (A20), anti-Ki67 (B56); PE-Cy7-conjugated anti-PD-1 (29F.1A12), anti-CD95 (Fas)(Jo2); PE-Dazzle 594-conjugated anti-B220 (RA3-6B2), anti-CXCR6 (SA051D1); BV421 anti-BCL6 (K112-91); BV605-conjugated anti-Ly-108 (SLAMF6) (13G3, BD Biosciences); BV650-conjugated anti-CD150 (SLAM)(TC15-12F12.2); BV711-conjugated anti-CXCR5 (L138D7); BUV395-conjugated anti-CD4 (GK1.5, BD Biosciences); FITC-conjugated anti-CD69 (H1.2F3); Unconjugated anti-TCF-1 (Cell Signaling Technology, C63D9).

Stained samples were analyzed with BD FACS LSR Fortessa, X20, or Symphony A3 or sorted on Aria II or III. Data were analyzed using FlowJo Software (FlowJo).

### Adoptive Transfer

For transfer into naive mice, CD4 T cells from the spleen and peripheral lymph nodes of C57BL/6 mice on 22-24 dpi with LCMV-c13 were harvested and enriched for CD4 T cells using a MojoSort Mouse CD4 T Cell Isolation Kit (BioLegend) prior to surface staining. 180,000 - 250, 000 Ly-108^−^ CXCR5^−^, Ly-108^+^ CXCR5^−^, or Ly-108^+^ CXCR5^+^ PD-1^+^ CD4 T cells were sorted and intravenously transferred into naive CD45.1 congenic recipients, which were infected with LCMV-c13 the next day. For infection-matched transfer, donor CD4 T cells from C57BL/6 mice on 21 dpi with LCMV-c13 were bead-enriched, surface stained and sorted as described above. 700,000 −1 million Ly-108^−^ CXCR5^−^, Ly-108^+^ CXCR5^−^, or Ly-108^+^ CXCR5^+^ PD-1^+^ CD4 T cells were sorted and transferred into infection-matched CD45.1 congenic recipients. Splenocytes from the recipient mice were harvested 15 d after transfer, and donor-derived CD4 T cells were analyzed by flow cytometry following magnetic bead–enrichment with MojoSort Mouse CD4 T Cell Isolation Kit (BioLegend).

### Tumor transplantation experiment

500,000 CFSE-labeled CD45.1/2 OT-II cells were intravenously transferred into C57BL/6 mice, which were subcutaneously inoculated with 1 million 1956-mOVA cells (Ferris et al., 2020) the next day. Tumor infiltrating lymphocytes (TILs) and cells in the tdLN were harvested 8 to 9 days after inoculation. LN cells were prepared by manual dissociation with frosted glass slides and TILs were prepared by digestion with Collagenase B, D1, and DNaseI, followed by staining with fluorescently labelled antibodies as described above.

### Statistical analysis

The *P* values were calculated with an unpaired two-tailed Student’s *t*-test and by one-way ANOVA for multigroup comparisons with the Tukey post hoc test using Prism 9 software (GraphPad): \**P* < 0.05, \*\**P* < 0.01, \*\*\**P* < 0.001, and \*\*\*\**P* < 0.0001.

