## Supplementary Figures for "BCL6-dependent TCF-1^+^ progenitor cells maintain effector and helper CD4 T cell responses to persistent antigen"

### Supplementary Materials

#### Supplementary Figure 1

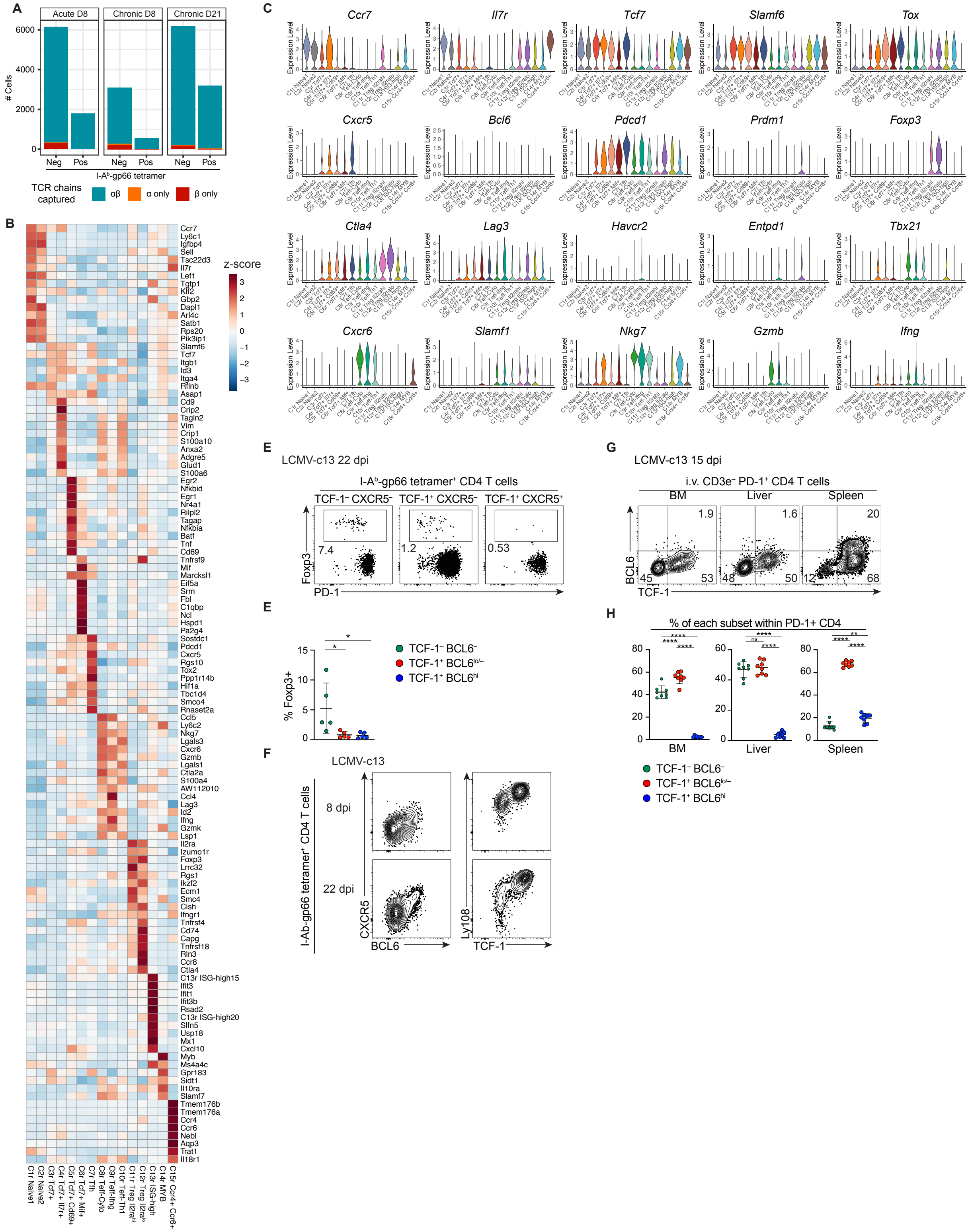

**Figure S1. scRNA-seq and flow analysis reveals heterogeneity within antigen-specific CD4 T cells in chronic LCMV infections. (Related to Figure 1)**
**(A)** TCRαβ chain capture efficiency of paired scRNA-seq and TCR-seq samples.
**(B)** Heatmap of the top 10 differentially expressed markers for each scRNA-seq cluster. Colors represent normalized expression per gene (z-score) among each cluster.
**(C)** Expression levels of select cluster markers and T cell subset defining genes within each cluster.
(**D, E**) Representative flow cytometry plots showing expression of PD-1 and Foxp3 in I-$A^b$-LCMV-gp66-specific splenic CD4 T cells in C57BL/6 mice infected with LCMV-c13 22 days before the analysis. Pooled data from 2 experiments with 2-3 mice / experiment are shown with mean±SD. Statistics done by one-way ANOVA.
**(F)** Representative flow plots showing correlation of CXCR5 vs BCL6 expression and Ly-108 vs. TCF-1 expression of I-$A^b$-LCMV-gp66 tetramer-specific splenic CD4 T cells in C57BL/6 mice infected with LCMV-c13 8 day or 22 days before the analysis.
**(G, H)** Representative flow plots showing expression of BCL6 and TCF-1 in PD-$1^+$ CD4 T cells in C57BL/6 mice infected with LCMV-c13 15 day before the analysis (G). Pooled data are shown in (H) with mean±SD. Statistics done with one-way ANOVA.

**Supplementary Figure 2**

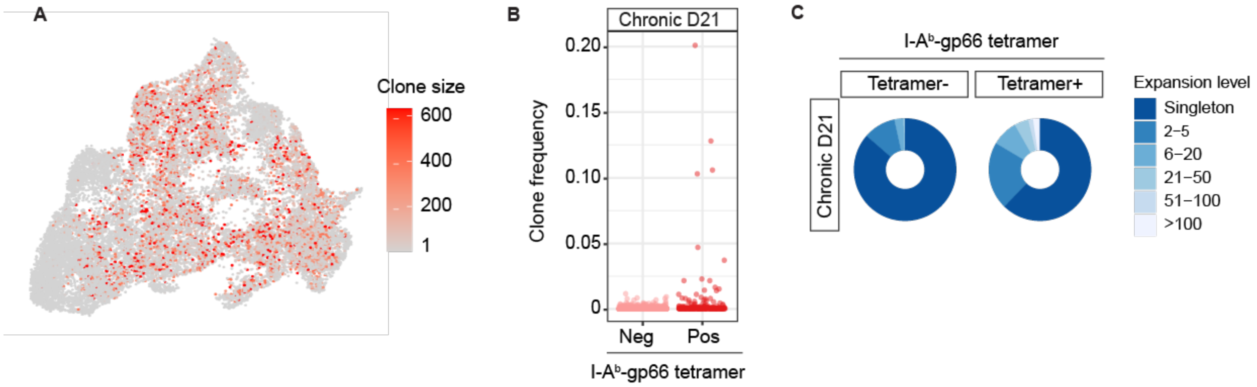


**Figure S2. Clonal expansion of I-A$^{b}$-gp66 Tet$^{+}$ T cells from LCMV-c13-21 dpi. (Related to Figure 2)**

**(A)** UMAP of LCMV-Arm 8 dpi, LCMV-c13 8 dpi and 21 dpi CD4 T splenocytes sorted for I-A$^{b}$-gp66 Tet$^{+}$ and Tet$^{-}$ populations, colored by clone size.

**(B)** Frequencies of TCR clones from I-A$^{b}$-gp66 Tet$^{+}$ and Tet$^{-}$ splenic CD4 T cells at 21 dpi with LCMV-c13.

**(C)** Clonal expansion levels of TCR clones from I-A$^{b}$-gp66 Tet$^{+}$ and Tet$^{-}$ splenic CD4 T cells at 21 dpi with LCMV-c13.

**Supplementary Figure 3**

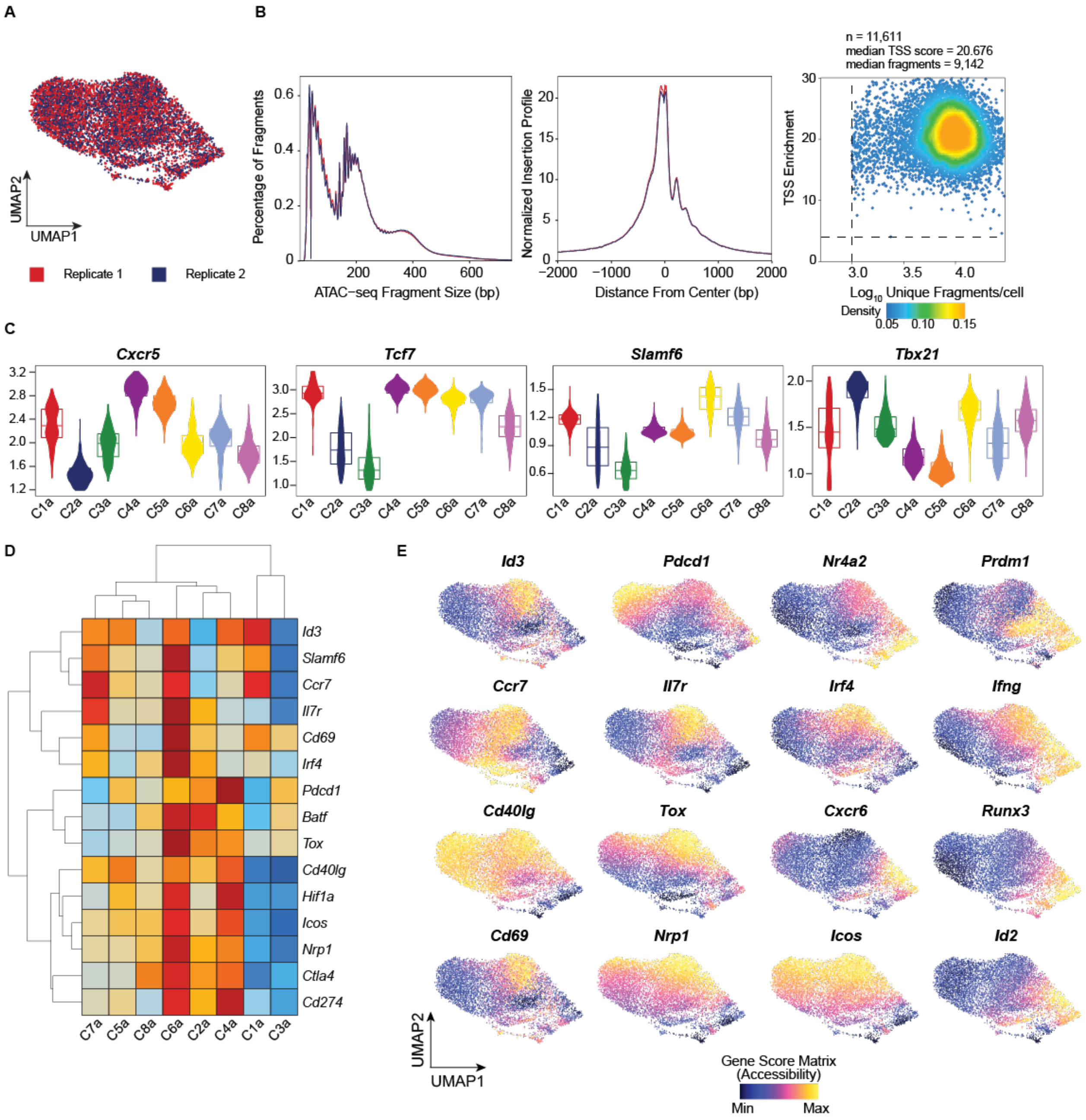


**Figure S3. scATAC-seq identifies chromatin features of Tprog, Tfh and Teff populations. (Related to Figure 3)**

**(A)** UMAP of scATAC-seq biological replicates of PD1$^{+}$ CD4 T cells sorted on 21 dpi of LCMV-cl13 infection.

**(B)** Quality control of scATAC-seq data. Fragment distribution of scATAC-seq libraries (left), Normalized insertion profile at the transcription start site (TSS) of genes (middle). Density plot of TSS read enrichment and unique fragments in single cells (right).

**(C)** Violin plot representation of gene scores for *Cxcr5*, *Tcf7*, *Slamf6* and *Tbx21*.

**(D)** Heatmap of marker gene accessibility scores of the progenitor population.

**(E)** UMAP of gene score (accessibility) values for the indicated genes.

**Supplementary Figure 4**

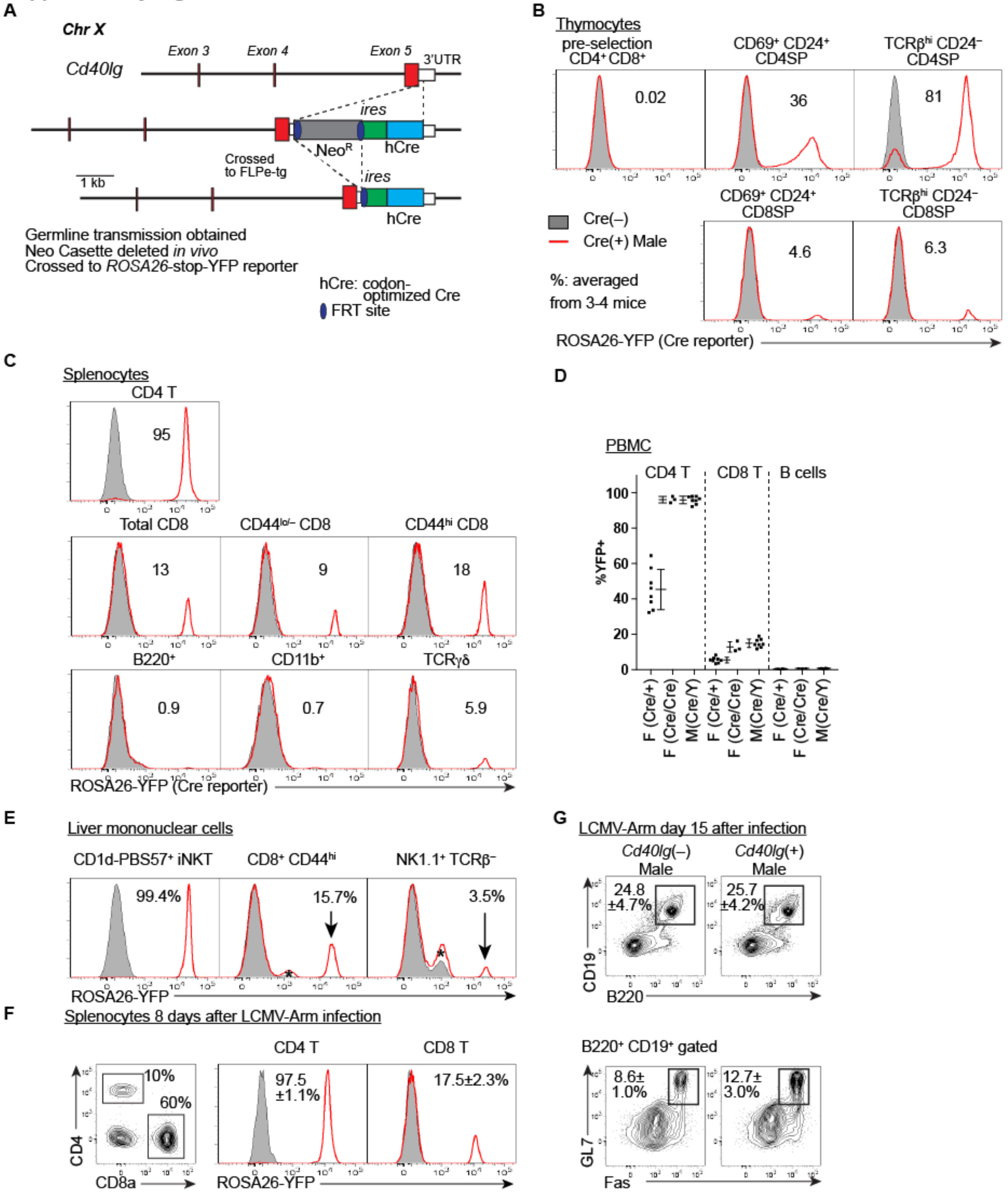


**Figure S4. The generation of *Cd40lg*-cre knock-in mice and validation of cre activity. (Related to Figure 5)**

**(A)** Targeted insertion of a codon-optimized cre (hCre) coding sequence followed by an internal ribosomal entry sequence into the 3' UTR of *Cd40lg* by homologous recombination in C57BL/6-derived JM8 ES cells. After obtaining germline transmission of the targeted allele, the FRT-flanked selection cassette was removed *in vivo* by crossing to a Flpe transgenic mice.

**(B-E)** Expression of YFP in thymocyte (B), splenocyte (C), peripheral blood mononuclear cell (D) and liver mononuclear cell (E) subpopulations from uninfected *CD40lg*-cre *Rosa26*(loxP-stop-loxP-YFP) mice. Data are shown with representative flow cytometry plots and mean±SD from 3-8 mice.
**(F, G)** Expression of YFP in splenic CD4 and CD8 T cells (F) and expression of GL7 and Fas by splenic B cells (G) in LCMV-Arm-infected *CD40lg*-cre *Rosa26*(loxP-stop-loxP-YFP) mice. Data are shown with representative flow cytometry plots and mean±SD from 3 mice. *P*-values by *t*-test: % of $B220^+$ $CD19^+$/spleen: $P = 0.815$, % of $GL7^+$ $Fas^+$/$B220^+$ $CD19^+$: $P = 0.122$.

**Supplementary Figure 5**

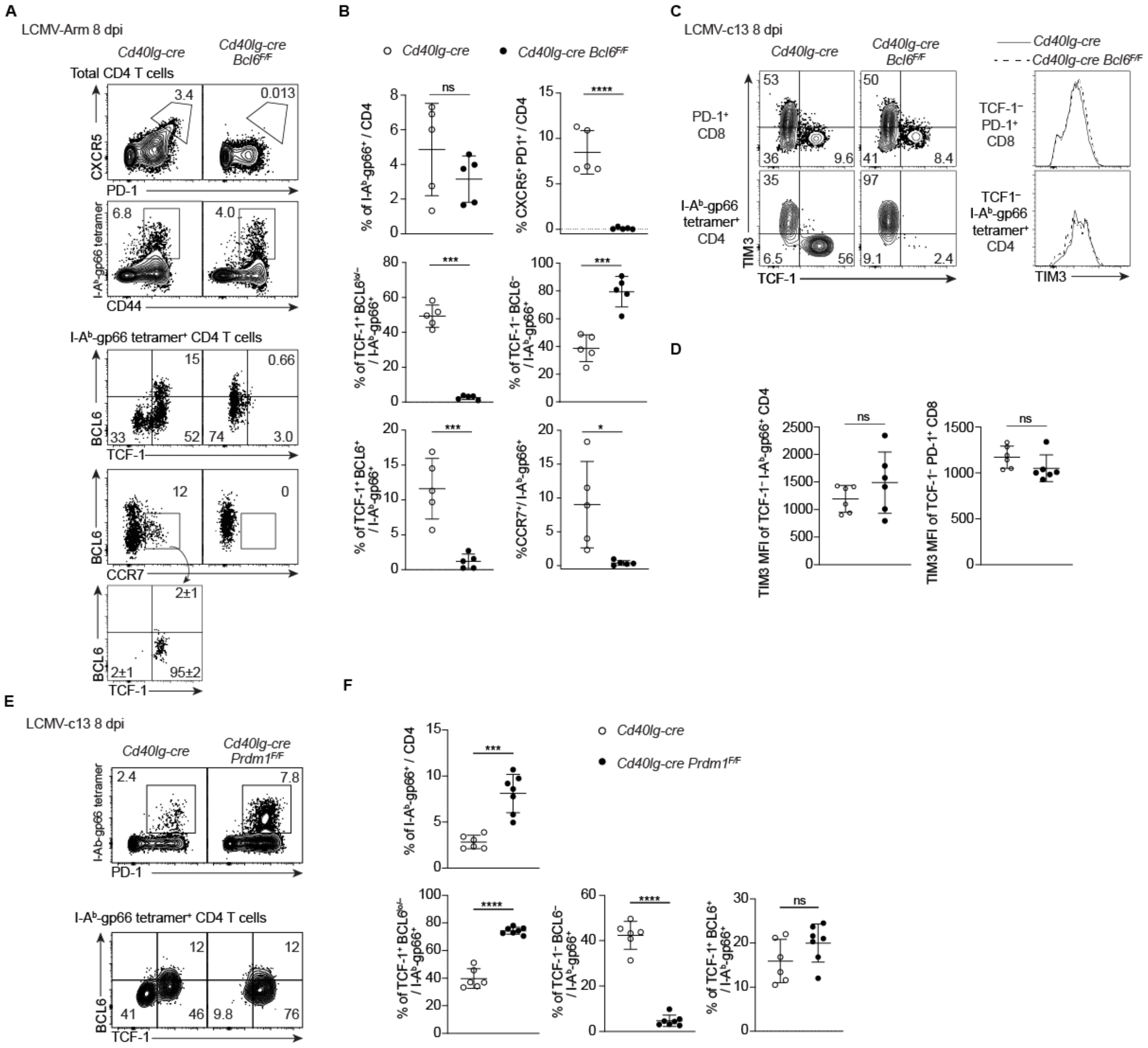


**Figure S5. BCL6 and Blimp1 are essential for the differentiation of TCF-1$^{+}$ and TCF-1$^{-}$ antigen-specific CD4 T cells, respectively, during LCMV infection. (Related to Fig 5)**

**(A, B)** Expression of PD-1, CXCR5, TCF-1, BCL6 and CCR7 in splenocytes from LCMV-Arm-infected *Cd40lg*-cre or *Cd40lg*-cre *Bcl6*$^{F/F}$ mice on 8 dpi. Representative flow plots (A) and pooled data from 2 experiments with n = 2-3 / genotype /experiment are shown with mean±SD in (B).

**(C, D)** Expression of TCF-1 and TIM3 in PD-1$^{+}$ CD8 T cells (C, top) and gp66-Tet$^{+}$ CD4 T cells (C, bottom) in the spleen of LCMV-c13-infected *Cd40lg*-cre or *Cd40lg*-cre *Bcl6*$^{F/F}$ mice on 8dpi. Histogram overlays of TCF-1$^{-}$-gated cells from each of the parental populations are shown (C, right). Mean fluorescence intensity (MFI) of TIM3 from 2 experiments with n = 2-4/genotype/experiment is shown with mean±SD in (D).

**(E, F)** Expression of TCF-1 and BCL6 by gp66-Tet$^{+}$ CD4 T cells in the spleen from LCMV-c13-infected *Cd40lg*-cre or *Cd40lg*-cre *Prdm1*$^{F/F}$ mice on 8 dpi. Representative flow plots (E) and pooled data from

2 experiments with n = 2-4 / genotype /experiment are shown in (F) with mean±SD. Statistical differences were assessed by unpaired *t*-test.
